# Spinal motoneuron excitability is homeostatically-regulated through β-adrenergic neuromodulation in wild-type and presymptomatic SOD1 mice

**DOI:** 10.1101/2024.03.25.586570

**Authors:** Stefano Antonucci, Guillaume Caron, Natalie Dikwella, Sruthi Sankari Krishnamurty, Anthony Harster, Hina Zarrin, Aboud Tahanis, Florian olde Heuvel, Simon M. Danner, Albert C. Ludolph, Kamil Grycz, Marcin Bączyk, Daniel Zytnicki, Francesco Roselli

## Abstract

Homeostatic feedback loops are essential to stabilize the activity of neurons and neuronal networks. It has been hypothesized that, in the context of Amyotrophic Lateral Sclerosis (ALS), an excessive gain in feedback loops might hyper- or hypo-excite motoneurons (MNs) and contribute to the pathogenesis. Here, we investigated how the neuromodulation of MN intrinsic properties is homeostatically controlled in presymptomatic adult SOD1(G93A) mice and in the age-matched control WT mice. First, we determined that β2 and β3-adrenergic receptors, which are Gs-coupled receptors and subject to tight and robust feedback loops, are specifically expressed in spinal MNs of both SOD1 and WT mice at P45. We then demonstrated that these receptors elicit a so-far overlooked neuromodulation of the firing and excitability properties of MNs. These electrical properties are homeostatically regulated following receptor engagement, which triggers ion channel transcriptional changes and downregulates those receptors. These homeostatic feedbacks are not dysregulated in presymptomatic SOD1 mice, and they set the MN excitability upon β-adrenergic neuromodulation.

## Introduction

Homeostatic plasticity is an essential mechanism that stabilizes neuronal network activity by regulating neuronal excitability (Turrigiano, 1999; Marder and Goaillard, 2006). Intrinsic excitability is set by a large panoply of ion channels (Smith and Brownstone, 2022) that are not only regulated at transcriptional level but are further subject to neuromodulation that controls their biophysical properties through post-translational modification (such as phosphorylation; Reckling *et al*., 2000). Indeed, neuromodulation acts through G protein-coupled membrane receptors (GPCRs) that activate or inactivate a number of intracellular kinases and G protein subunits targeting ion channels (Carr *et al*., 2003; Luo *et al*., 2022; Gonzalez-Hernandez, 2024). However, signaling of the GPCRs itself is subject to multiple feedback loops. A well established example is the desensitization of the β-adrenergic receptors, mainly coupled to the Gs protein, during prolonged exposure to their agonists (Maaliki *et al.,* 2024). Their desensitization can be caused by several mechanisms involving surface localization, protein stability and transcriptional downregulation and mRNA stability (Bouvier *et al*., 1989; Okeke *et al.,* 2019). Upon activation of the receptor, the Gs protein activates the cAMP/PKA pathway, inducing phosphorylation of the GPCR itself and, as a consequence, the recruitment of β-arrestins on the adrenoceptor and the endocytosis of the complex (Maaliki *et al*., 2024). Prolonged agonist exposure can also induce a decreased transcription or mRNA stability, reducing mRNA abundance (Bouvier *et al*., 1989). Receptor desensitization results in a reduced neuromodulation, which represents a short-term feedback that contributes to overall homeostatic plasticity.

In Amyotrophic Lateral Sclerosis (ALS), vulnerable spinal motoneurons (MNs) and the neuronal networks that control their firing display abnormalities from early motor presymptomatic phases onwards (Bączyk *et al*., 2022). Indeed, in the SOD1(G93A) ALS murine model (henceforth, SOD1), alterations of intrinsic electrical properties of MNs and synaptic inputs to MNs are evolving from embryonic to juvenile and adult time points (Quinlan *et al*., 2011; Martin *et al*., 2013; Branchereau *et al*., 2019; Leroy *et al*., 2014; Delestrée *et al*., 2014; Martínez-Silva *et al*., 2018; Branchereau *et al*., 2019; Bączyk *et al*., 2020; Jensen *et al*., 2020; Cavarsan *et al*., 2023; Nascimento *et al*., 2024). While it has been hypothesized that an excessive gain in homeostatic feedback loops (Brownstone and Lancelin, 2018; Kuo *et al*., 2020) might hyper- or hypo-excite MNs and drive the pathogenesis of ALS, the occurrence of actual alterations of homeostatic responses of ALS MNs has not been demonstrated.

To test this hypothesis, we investigated if the neuromodulation of MN intrinsic properties is homeostatically controlled in presymptomatic adult SOD1 mice and age-matched wild-type (WT) controls. First, we determined which Gs-coupled receptors are specifically expressed in MNs and are not dysregulated in adult presymptomatic SOD1 mice. Among them, we found β2 and β3- adrenergic receptors, which are known to be subject to tight and robust feedback loops. We then studied *in vivo* how these receptors increased MN firing, through cAMP/PKA activation, both in WT and SOD1 MNs, showing a so far overlooked neuromodulation of MNs through β-adrenergic receptors. Most importantly, we demonstrated that, upon agonist-driven chronic activation, both β2 and β3 receptors underwent homeostatic downregulation, desensitizing MNs to the β-adrenergic neuromodulation. This homeostatic response is the same in MNs from SOD1 and WT mice, indicating that this form of homeostatic plasticity is maintained at a presymptomatic disease stage.

## Results

### 1. Expression of β-adrenergic receptors in MNs of WT and ALS presymptomatic mice

Since β-adrenergic receptors, as well as multiple other PKA-coupled GPCRs, might be subject to different degrees of disease-related modulation and disease-associated transcriptional changes, we first set out to verify their pattern of expression at P45 (presymptomatic stage; Martínez-Silva *et al*., 2018; Bączyk *et al*., 2020) in lumbar MNs from SOD1 and WT animals isolated by laser-capture-microdissection (LCM; 30 MNs > 500μm^2^ from L4-L5, as in Song *et al*., 2023) (Fig. 1A). Notably, the β-adrenergic receptors (*Adrb1-3*) were highly expressed in MNs and either upregulated (*Adrb1*) or showing minimal downregulation (*Adrb2*) or unmodified expression (*Adrb3*) in ALS MNs. We then focused on this receptor family because of their well-known desensitization property upon prolonged agonist exposure and the lack of knowledge of their physiological role in MNs.

**Figure 1.**
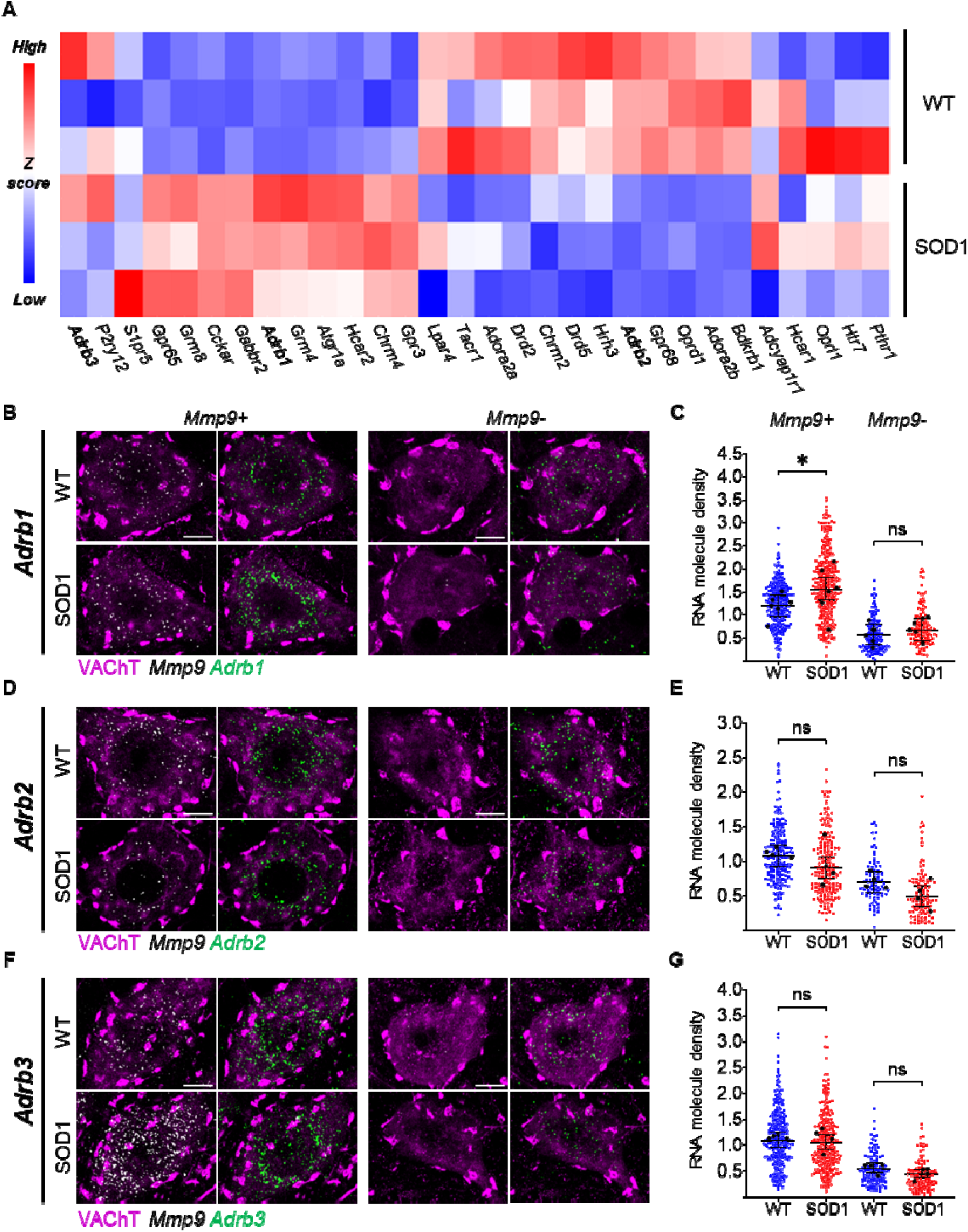
Adrenergic β2 and β3 receptors are expressed in WT and ALS motoneurons and minimally affected at presymptomatic stage. **A)** GPCR-targeted transcriptomics in laser-microdissected MNs reveals that 9/30 GPCRs are downregulated in SOD1 MNs (*Oprd1*, *Adora2b*, *Bdkrb1*, *Drd5*, *Chrm2*, *Adrb2*, *Gpr68, Drd2, Hrh3),* 10/30 are upregulated (*Gpr65*, *Grm8*, *Cckar*, *Gabbr2*, *Adrb1*, *Grm4*, *Hcar2*, *Atgr1a*, *Gpr3* and *Chrm4*) and 11/30 GPCR (*Adcyap1r1, Hcar1*, *Oprl1*, *Pthr1*, *Htr7*, *Adrb3*, *P2ry12*, *Lpar4*, *S1pr5, Tacr1* and *Adora2a*) are comparably expressed. Of note, *Adrb2* and *Adrb3* are minimally affected (*blue*, downregulated genes; *red*, upregulated genes). 30 MNs were pooled per animal; N = 3 animals per group. **B-C)** Single-molecule *in situ* hybridization (ISH) confirms the upregulation of *Adrb1* mRNA in *Mmp9*+ (vulnerable), but not in *Mmp9*- (resistant) MNs. **D-G)** ISH reveals the comparable expression of *Adrb2* (D-E) and *Adrb3* (F-G) in WT and SOD1 MNs. All confocal images show single representative MNs (left panel, merged VAChT and *Mmp9* mRNA; right panel, merged VAChT and adrenoceptor mRNA). Scale bar = 10µm. Dotplots quantify individual cell expression levels as the number of mRNA molecules per µm^2^. Small dots are individual MN data points and large black dots indicate average expression per animal; bars represent mean and 95% confidence intervals. N = 4-7 mice per group. \**p*<0.05, ns - non significant.

We validated the expression of *Adrb1-3* by single-molecule *in situ* hybridization (ISH) in lumbar MNs, distinguishing MN subtypes according to *Mmp9* expression (*Mmp9*+ MNs are disease-vulnerable innervating Fast-contracting Fatigable (FF) and large Fast-contracting Fatigue-Resistant (FR) motor units, whereas *Mmp9*- MNs are disease-resistant innervating small FR and Slow-contracting (S) motor units; Kanning *et al*., 2010; Kaplan *et al*., 2014; Leroy *et al*., 2014). All MNs variably express *Adrb1-3* mRNAs. However *Adrb1-3* mRNA was more abundant in *Mmp9+* than in *Mmp9-* MNs (50% or fewer mRNA molecules in *Mmp9-* MNs; Fig. 1B-G). Nevertheless, the disease-related modulation was largely (but not completely) similar in the two subpopulations: *Adrb1* was upregulated only in *Mmp9*+ MNs (Fig. 1B-C), whereas *Adrb2* (Fig. 1D-E) and *Adrb3* (Fig. 1F-G) were not significantly affected in either subpopulation (all statistics are reported in Supplemental Excel Table 1). The β2 and β3 receptors are then amenable to be probed in the context of their homeostatic loops since their genes expression is unaffected by the disease process at P45.

### 2. A punctual β2 and β3 agonists delivery activates PKA signaling and expression of immediate-early genes

Prior to their use to study β-adrenergic desensitization feedback in MN, we demonstrated that two FDA-approved adrenergic β2 and β3 agonists (formoterol and amibegron, respectively) cross the blood-brain barrier and engage signal transduction in MN, and can elicit a transcriptome modulation in MNs. We de-prioritized the β1 receptor because it was already affected by the disease, agonists do not cross the blood-brain barrier well and they produce substantial systemic effects. WT or SOD1 mice were administered β2/β3 agonists (noted AF in all figures) or vehicle, and PKA cascade activation was assessed at 3h in spinal cord sections by immunolabeling for the PKA-phosphorylation consensus motif RRxpS/T. In vehicle-treated SOD1 animals (baseline PKA activity), MNs displayed a lower RRxpS/T immunostaining than WT (Fig. 2A). However, agonist treatment elicits a substantial increase in RRxpS/T immunoreactivity (Fig. 2A), indicating target engagement (increased in RRxpS/T immunoreactivity was also observed in vasculature; Fig. 2A and Supplemental Excel Table 1). Coherent with the signaling engagement, in LCM-dissected MNs, β2/β3 adrenergic agonists upregulated the expression of multiple immediate-early genes both in WT and SOD1 animals (3h after treatment; in particular, *cFos*, Δ*FosB*, *Npas4* and *Egr1* were induced in WT and *Fos*, Δ*FosB* only in SOD1 animals; Fig. 2B and Supplemental Excel Table 1). Taken together, these data demonstrate that β2/β3 receptors on murine MNs are engaged by agonists and linked to signaling and transcriptional responses.

**Figure 2.**
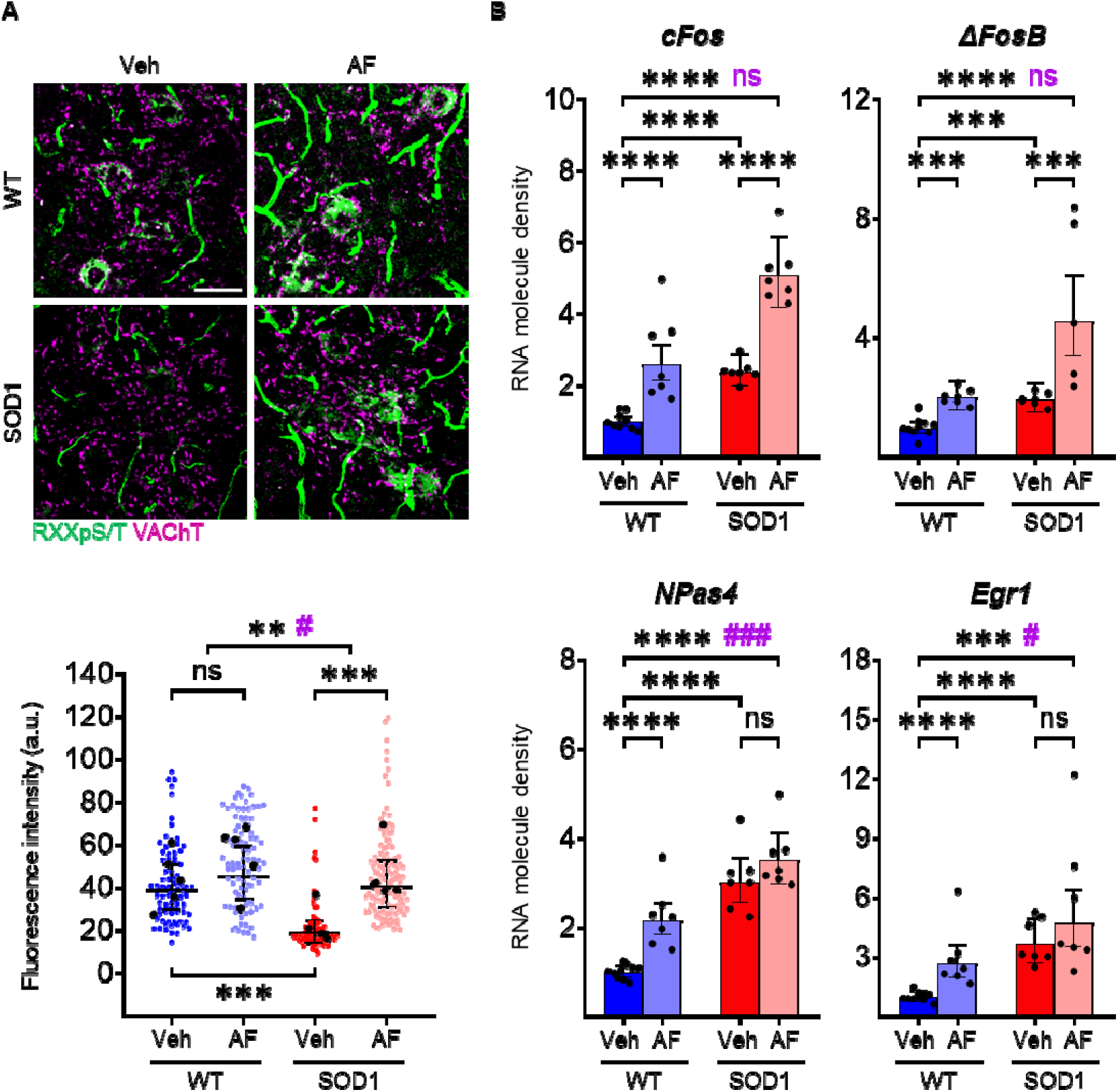
Punctual pharmacological activation of β2/β3 receptors activates PKA signaling and induces Immediate Early Gene transcription in motoneurons. **A)** Immunofluorescence reactivity for the consensus PKA-phosphorylated sequence RRxpS/T reveals increased immunoreactivity in the cytoplasm of MNs and in spinal vasculature upon β2/β3 stimulation in WT (strong trend, significant when the outlier animal is excluded) and SOD1 animals (scale bar = 50µm). In scatterplots, small dots correspond to individual MN data points, large black dots to average per animal; mean and 95% confidence intervals are depicted. N = 5 animals per group. Significances on top bars illustrate treatment effects across genotypes (WT and SOD1 grouped together; black) or interaction effects (magenta). *Post-hoc* significances are shown for WT Veh *vs.* WT AF (treatment effect in WT), SOD1 Veh *vs.* SOD1 AF (treatment effect in SOD1) and for WT Veh *vs.* SOD1 Veh (genotype effect before treatment, bottom bar). **B)** Induction of immediate-early gene mRNAs (*cFos*, Δ*FosB*, *NPas4*, *Egr1*) in MNs upon β2/β3 stimulation. Each dot represents an individual animal. Significance bars illustrate treatment effects across genotypes (WT and SOD1 grouped together; black) or interaction effects (magenta). *Post-hoc* significances are shown for WT Veh *vs.* WT AF (treatment effect in WT, lower left bar), SOD1 Veh *vs.* SOD1 AF (treatment effect in SOD1, lower right bar) and for WT Veh *vs.* SOD1 Veh (genotype effect before treatment, middle bar). N = 7-10 animals per group. *^/#^*p*<0.05, \*\**p*<0.01, ***^/###^*p*<0.001, \*\*\*\**p*<0.0001, ns - non significant.

### 3. A punctual adrenergic β2/β3 activation increases MN excitability in both WT and SOD1 mice

The neuromodulatory properties on MNs of the β2 and β3-adrenergic receptors are unknown so far. Then, before investigating the homeostatic responses triggered by a long exposure to agonists we needed to elucidate how intrinsic electrical properties of MNs are modulated by a punctual delivery of the agonists. Our physiological studies relied on *in vivo* intracellular recordings carried out on anesthetized WT and SOD1 mice. To investigate the neuromodulatory properties, we first recorded, in each experiment, a sample of hindlimb MNs (between 1 and 7), then we intravenously (i.v.) delivered a single bolus of the adrenergic β2/β3 agonists (AF) before recording a new sample of MNs (between 1 and 9) in the next 3 hours following the drug delivery (see Methods section).

A series of depolarizing or hyperpolarizing pulses of currents (Fig. 3A-B) and a triangular slow ramp of current (Fig. 4A-B) allowed us to investigate whether the treatment with adrenergic β2/β3 agonists modify the physiological parameters that determine MN excitability (resting membrane potential, input conductance, voltage threshold for firing, recruitment current, firing gain; all statistics are detailed on Supplemental Excel Table 1). We found that the treatment does not significantly change the resting membrane potential both in WT and SOD1 MNs (Fig. 4C), nor the input conductance measured on the response peak (Gin Peak) at the onset of current pulses (Fig. 3C). In response to long pulses, the peak is followed by a sag down to a plateau that is caused by an H-current (Fig. 3A) (Ito and Oshima, 1965; Takahashi, 1990; McLarnon, 1995; Manuel *et al*., 2009). We then also measured the input conductance at the end of the plateau (Gin Plateau, Fig. 3A-B) and we found that the treatment significantly reduces Gin plateau (Fig. 3D) across genotypes (WT and SOD1) (Fig. 3E) suggesting that H-current has decreased. An almost significant interaction effect in Gin Plateau suggests differences in treatment efficacy between genotypes with a stronger effect in SOD1 mice. In the same line, the input conductance measured at the beginning of the slow ramp (Gin ramp, Fig. 4B) is differently affected by treatment between WT and SOD1 mice (Fig. 4D). The treatment causes a significant reduction of Gin ramp in SOD1 mice but not in WT mice. It then appears that β agonists decrease Gin plateau and Gin ramp in MNs from SOD1 mice, likely through a modulation of H-current. This effect might increase MN excitability during slow depolarizing events.

**Figure 3.**
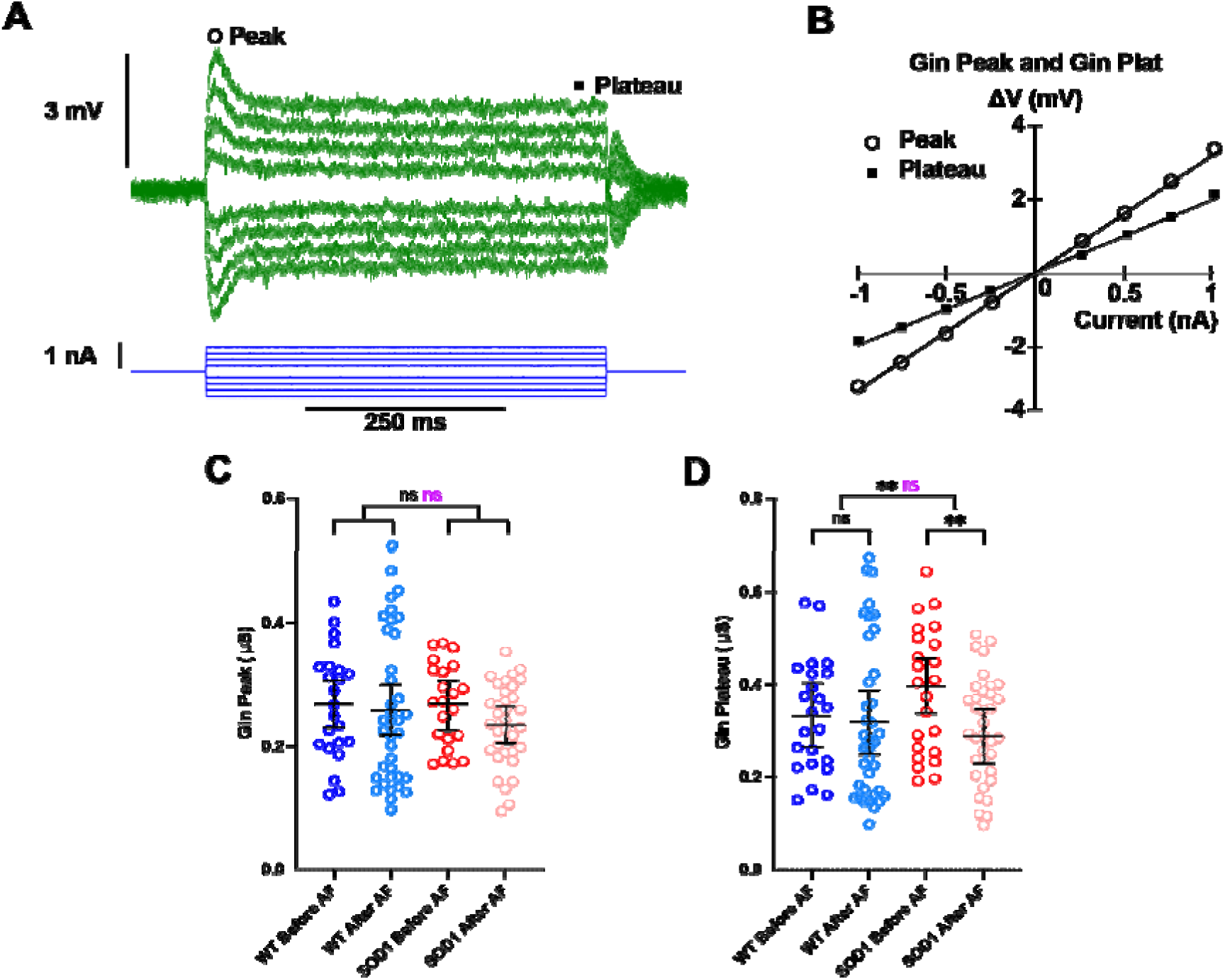
Punctual delivery of adrenergic β2/β3 agonists decreases the plateau input conductance of motoneurons in SOD1 mice. **A)** Average responses of a SOD1 MN to a series of current pulses lasting 500ms and ranging from −1 to +1nA. Notice that, in each voltage response, the peak is followed by a sag before it stabilizes to a plateau. **B)** Plot of the voltage-current response measured at peak and plateau against the intensity of current. Effect of β2/β3 agonists on peak input conductance **(C)** and plateau input conductance **(D)** in MNs from WT and SOD1 mice. Significances on top bars are for treatment effects (black) and interaction effects (magenta). *Post-hoc* significances are shown for WT before AF *vs.* WT after AF (treatment effect in WT), SOD1 before AF *vs.* SOD1 after AF (treatment effect in SOD1) and for WT before AF *vs.* SOD1 before AF (genotype effect before treatment, bottom bar). Amibegron + Formoterol (AF). N = 6 WT mice and N = 11 SOD1 mice. \*\**p*<0.01, ns - non significant.

**Figure 4.**
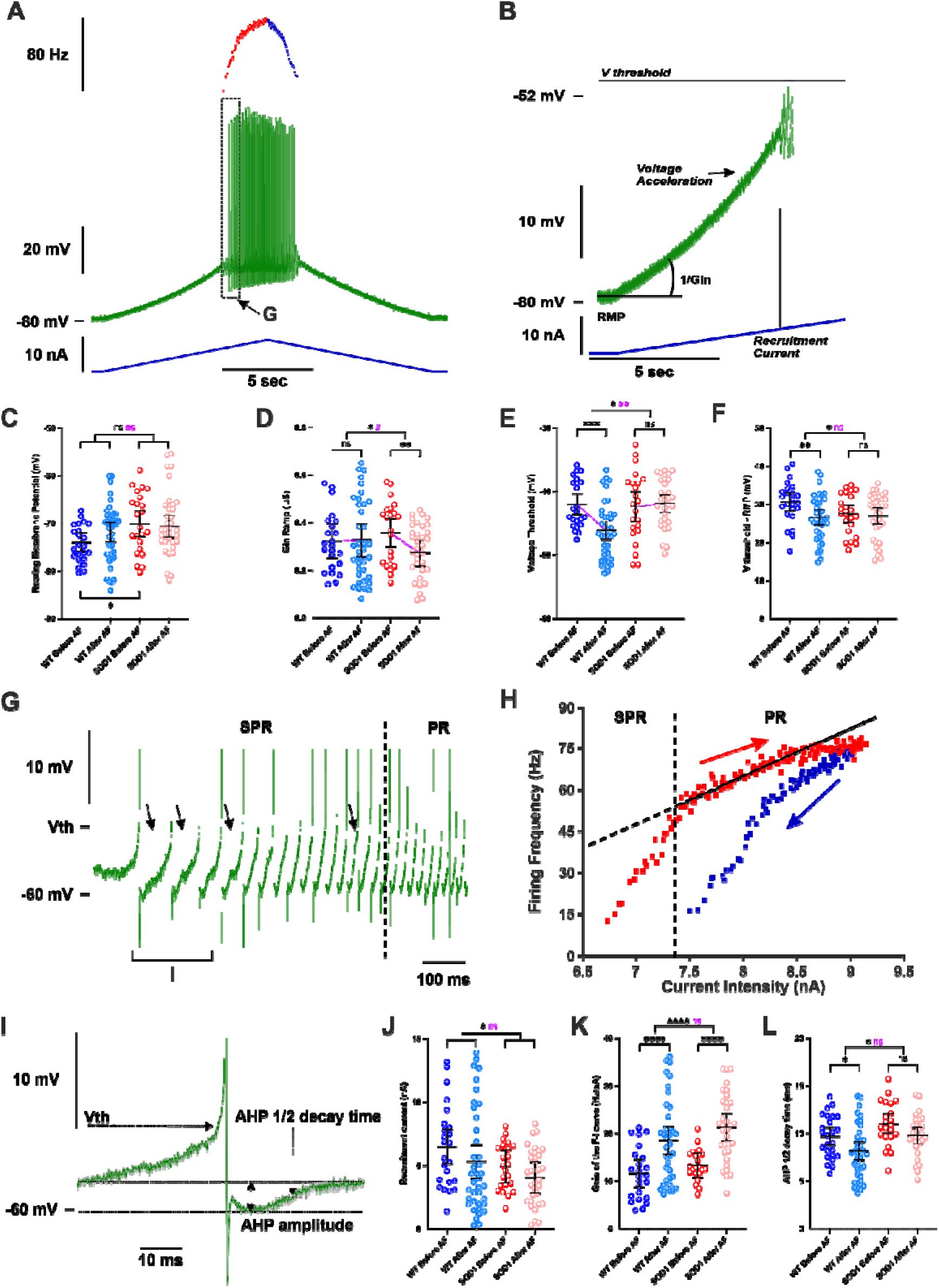
Punctual delivery of adrenergic β2/β3 agonists increases the firing of motoneurons in WT and SOD1 mice. **A)** Representative response of a SOD1 MNs to a slow ramp (1nA/s) of current. Ramp of current (blue bottom trace), voltage response (green middle trace) and instantaneous firing frequency (top trace with ascending leg in red and descending leg in blue). **B)** Magnification of the ramp subthreshold voltage (green top trace) and current (blue bottom trace). Notice that Vthreshold (Vth) was measured in expanded traces as shown in **G** and **I**. Effect of an acute delivery of Amibegron/Formoterol (AF) on resting membrane potential **(C)**, ramp input conductance **(D)**, voltage threshold for spiking **(E)**, voltage threshold - resting membrane potential **(F)**, recruitment current **(J)**, Gain of the F-I relationship **(K)**, AHP half-decay time **(L)** in MNs from WT and SOD1 mice. **G)** Magnification and time expansion of the voltage trace from the region indicated in **A**. At the firing onset, oscillations (arrowheads) appear in the interspike intervals, which characterizes the subprimary range (SPR). When the injected current increases oscillations disappear, reducing the firing variability and increasing the firing frequency linearly, which characterizes the primary range (PR). **H)** Plot of the instantaneous firing frequency against the intensity of current for the ascending (red) and descending (blue) legs of the ramp shown in **A**. The F-I relationship displayed a clockwise hysteresis. Vertical dashed line indicates the transition between the SPR and the PR on the ascending leg. The gain of the F–I curve can be estimated by the slope of the linear regression (continuous line) in the PR. **I)** Average of the three first action potentials displayed in **G.** The AHP half-decay time is the time between AHP peak amplitude and where the AHP relaxed to half its amplitude. In all graphs, each point represents one MN and the mean ± 95% confidence intervals are shown. Significances on top bars are for treatment effects across genotypes (WT and SOD1 grouped together; black) or interaction effects (magenta). *Post-hoc* significances are shown for WT before AF *vs.* WT after AF (treatment effect in WT), SOD1 before AF *vs.* SOD1 after AF (treatment effect in SOD1) and for WT before AF *vs.* SOD1 before AF (genotype effect before treatment, bottom bar). Amibegron + Formoterol (AF). N = 6 WT mice and N = 11 SOD1 mice. *^/#^*p*<0.05, **^/##^*p*<0.01, \*\*\**p*<0.001, \*\*\*\**p*<0.0001, ns - non significant.

Another parameter that sets MN excitability is the voltage threshold at which spikes are fired (Vthreshold): the more hyperpolarized the voltage threshold for spiking, higher is the probability to elicit firing. We found that treatment causes a significant hyperpolarization of the voltage threshold in WT mice, suggesting an increased excitability, but not in SOD1 mice (Fig. 4E). The difference between voltage threshold and RMP (Vthreshold−RMP) is indeed significantly smaller after treatment compared to before across genotypes (Fig. 4F). However, while post-hoc tests showed that the treatment only had a significant effect in WT and not in SOD1 mice, the difference of the treatment effect between the two genotypes was not significant for Vthreshold−RMP. The next important parameter for assessing excitability is the amount of depolarizing current required to reach the firing threshold, *i.e.* the recruitment current. During slow ramps, the voltage trajectory to reach the firing threshold is not linear since an acceleration occurred below the threshold for firing (Fig. 4B, arrow). We found that treatment has a slight effect on the recruitment current which is reduced across genotypes (WT and SOD1 mice) (Fig. 4J), contributing to the excitability increase.

Once the discharge is initiated, the MN starts to discharge in the mixed mode oscillations regime, in which small oscillations (arrows in Fig. 4G) are present between spikes, describing a subprimary range, before transitioning to a regime of firing with full blown spikes whose frequency increases linearly with the injected current, *i.e.* the primary range (Fig. 4G, and see Manuel *et al*., 2009). Another important criteria to assess MN excitability is the slope (frequency-current gain) of the primary firing range (Fig. 4H), with a higher frequency-current intensity gain (F-I gain) indicating that the MN is more excitable. Indeed, the strongest action of β2/β3 agonists was observed on the F-I gain that is very significantly increased in MNs from both WT (64% increase) and SOD1 mice (63% increase) (Fig. 4K). Then, we asked, whether a faster kinetics of AHP current might contribute to the increase of the F-I gain (Meunier and Borejsza, 2005; Manuel *et al*., 2006). We systematically measured the half-relaxation decay time on the AHP that follows the first 3-5 averaged spikes in the subprimary range, a range where AHP fully relaxed before the next spike (see Methods section; Fig. 4I). Interestingly, the treatment caused significant shortening of the AHP half decay time across genotypes (Fig. 4L). The faster AHP kinetics is likely contributing to the gain increase.

To sum up, an acute administration of β2/β3 agonists increases MN excitability mainly through an increase of the F-I gain in both WT and SOD1 mice. However, a decrease of Gin plateau or Gin ramp in MNs from SOD1 mice, and a hyperpolarization of voltage threshold for spiking in MNs from WT mice may also contribute to the excitability increase induced by β2/β3 agonists.

### 4. A punctual and direct injection of cAMP analogue also increases MN excitability in SOD1 mice

To confirm that the increase in the F-I gain of MNs upon a punctual activation of the β2/β3 adrenergic receptors was mediated by the cAMP/PKA pathway (and not to alternative signaling, such as β-arrestin/ERK; Huang *et al*., 2018) and was not an off target effect (*i.e*. increase in heart rate, etc…), we directly activated the cAMP/PKA pathway by intracellular injection in MN of the cAMP analogue (cAMP-SP; Bączyk *et al*., 2020). We first recorded, before the injection, the responses of the MNs to a slow ramp of current and to long pulses (in order to determine the peak and the plateau input conductances). Next, we electrophoretically injected cAMP-SP through the intracellular microelectrode (normalized electrical load 2600 ± 600nA.sec/µS). We then recorded again the MN responses to slow ramps of currents and long pulses during at least 10 minutes. The whole procedure required that the MN recording did not deteriorate for at least 30 minutes, a condition that limited the number of successful experiments (see Methods section).

We have been able to follow the effect of cAMP-SP in 7 MNs from SOD1 mice (Fig. 5, statistics in Supplemental Excel Table 1). cAMP-SP significantly hyperpolarized the resting membrane potential (Fig. 5C), and it slightly decreased the input conductance during the ramp (Gin Ramp, Fig. 5D) (but it had no effect on the peak input conductance and plateau input conductance during long pulses). cAMP-SP also hyperpolarized the voltage threshold for spiking (Fig. 5E) but it did not significantly modify Vthreshold−RMP (Fig. 5F), nor the recruitment current (Fig. 5G).

**Figure 5.**
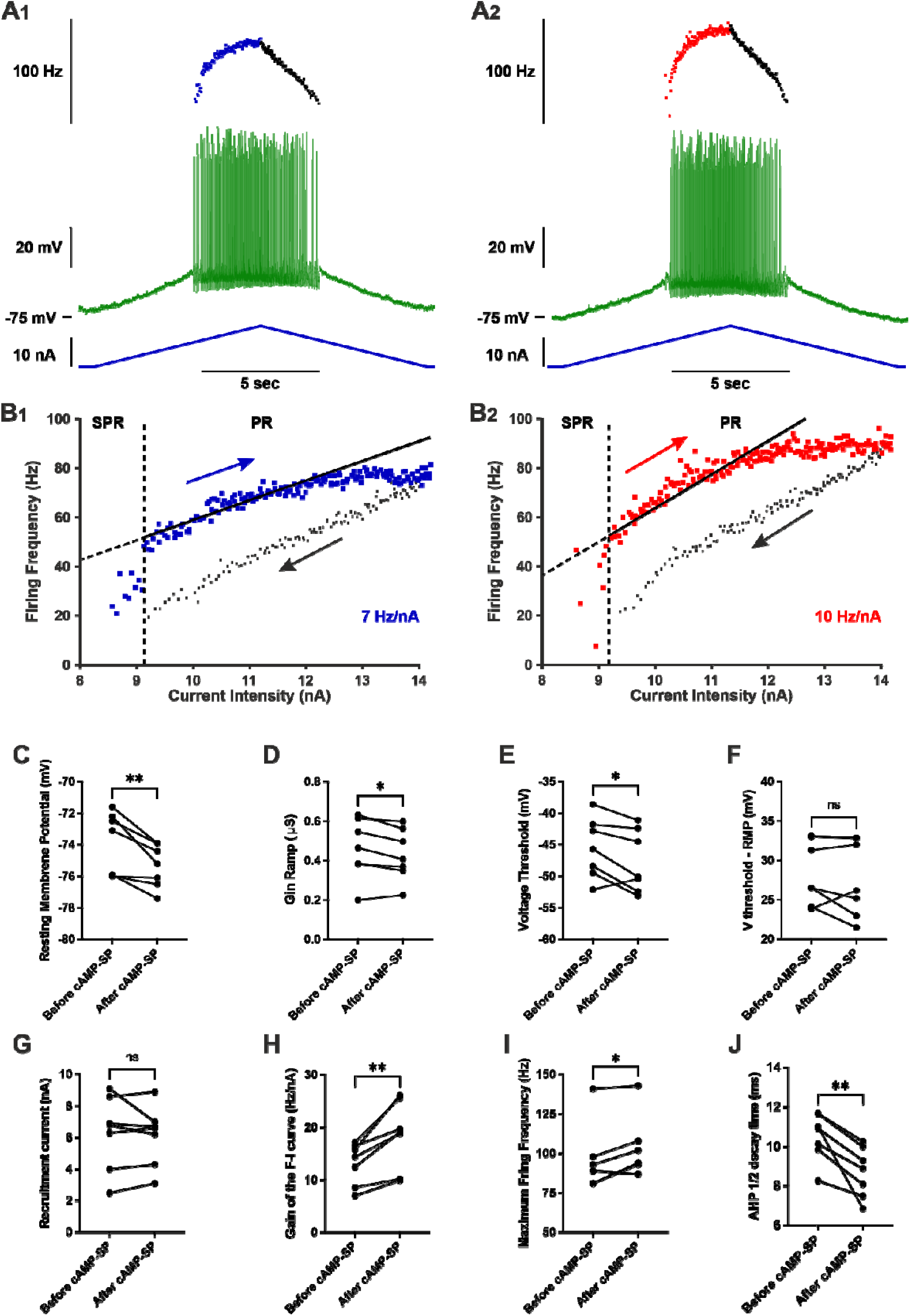
Punctual and direct activation of cAMP/PKA pathway increases the firing of motoneurons in SOD1 mice. **A)** Representative response of a SOD1 MNs to a slow ramp (2nA/s) of current, before (**A_1_**) and 12 minutes after (**A_2_**) the iontophoretic injection of cAMP-SP in the same instantaneous firing frequency (top trace with ascending leg in blue for **A_1_** or red for **A_2_**, and descending leg in black for both **A_1_** and **A_2_**. **B)** Plot of the instantaneous firing frequency against the intensity of current for the ascending (blue for **B_1_** or red for **B_2_**) and descending legs (black for both **B_1_** and **B_2_**) of the ramps shown in **A**. The F-I relationships displayed a clockwise hysteresis. Vertical dashed line indicates the transition between the SPR and the PR on the ascending branch. The gain of the F–I relationship can be estimated by the slope of the linear regression (continuous line) in the PR. **C-J)** Comparison of electrophysiological properties extracted from the slow ramp of current before and after injection of cAMP-SP. Before values are measured just before the injection; After values are averages of the recordings repeated after the iontophoretic injection of cAMP-SP; n=7 MNs recorded in N = 7 SOD1 mice. **C)** Resting membrane potential. **D)** Ramp input conductance. **E)** Voltage threshold for spiking. **F)** Voltage threshold - resting membrane potential. **G)** Recruitment current. **H)** Gain of the F-I curve. **I)** Maximum firing frequency reached at the end of the ascending ramp (the velocity and amplitude were the same for all ramps in each MNs). **J)** AHP half-decay time. In all graphs, each linked two points represent one MN. Before *vs*. after values were compared using a paired *t*-test except for maximum firing frequency, for which a Wilcoxon paired test was used. \**p*<0.05, \*\**p*<0.01, ns - non significant.

However cAMP-SP significantly increased the F-I gain (average increase 32%, Fig. 5H) and, consequently, the maximal firing frequency measured at the end of the ramps (ramp velocity and amplitude were kept constant before and after cAMP-SP injection for each individual motoneuron) (Fig. 5I). In addition, cAMP-SP shortened the AHP half decay time as acute β2/β3 agonists did (Fig. 5J). Altogether, many actions elicited in MNs by a direct activation of the PKA pathway (decrease of Gin Ramp, hyperpolarization of the voltage threshold, increase of the F-I gain, faster AHP kinetics) are the same as those produced by an acute i.v. delivery of β2/β3 agonists. These results strongly suggest that most of the β2/β3 agonists effects are caused by the activation of the cAMP/PKA pathway in MNs.

### 5. A punctual β2 and β3 agonists delivery is sufficient to induce a dysregulation of ion channels transcription

Since β2/β3 adrenergic agonists exerted a transcriptional modulation already with 3h of administration (as shown by the induction of immediate-early genes), we explored whether they would also influence the genes encoding for ion channels responsible for MN excitability. We investigated the expression of 19 ion channel-encoding genes responsible for K^+^ (*Kcna1, Kcna2, Kcnab1, Kcnb1, Kcnj14*, *Kcnn2*, *Kcnn3*, *Kcnq2*, *Kcnq3*, *Kcnq5*, *Kcnt2*), Na^+^ (*Scn1a*, *Scn8a*), Ca^2+^ (*Cacna1d*, *Cacna2d3*), Cl^-^ (*Tmem16f*) and non-selective cationic currents (*Hcn1, Hcn2, Trpm5*). A large fraction of the channels under analysis exhibited significant disease-driven changes already at this presymptomatic stage (14/19, with only *Cacna2d3*, *Hcn2*, *Kcnq2*, *Scn8a* and *Tmem16f* remaining unaffected, see Supplemental Excel Table 1). Remarkably, three hours after β2/β3 agonist administration, the majority of investigated channels was further significantly modulated (down- or up-regulated) not only in WT but also in SOD1 MNs (Fig. 6A), with a clear distinction of genotypes and treatment groups upon principle component analysis (Fig. 6B). Specifically, upon treatment both WT and SOD1 MNs displayed a shared pattern of downregulation of Ca^2+^ (*Cacna2d3*), Na^+^ (*Scn1a*), K^+^ (*Kcna1, Kcna2, Kcnab1*), mixed cationic (*Hcn1, Hcn2, Trpm5*) or Cl^-^ (*Tmem16f*) channels and upregulation of other K^+^ channels (*Kcnq3, Kcnq5*). Overall, 11 out of 19 investigated genes showed similar modulation of their transcriptional responses in AF-treated WT and SOD1 when compared to vehicle-treated littermates (for detailed statistics, see Supplemental Excel Table 1). *Cacna1d*, *Kcnb1*, *Kcnj14*, *Kcnn3*, *Kcnq2* and *Scn8a* were not significantly affected by the adrenergic agonist treatment in both genotypes whereas only two, *Kcnn2* and *Kcnt2,* showed significant changes in opposite directions for WT and SOD1 genotypes. These data demonstrate that a punctual stimulation of β2/β3 receptors is sufficient to elicit a rapid modulation of the MN channelome that is similar in WT and SOD1 animals.

**Figure 6.**
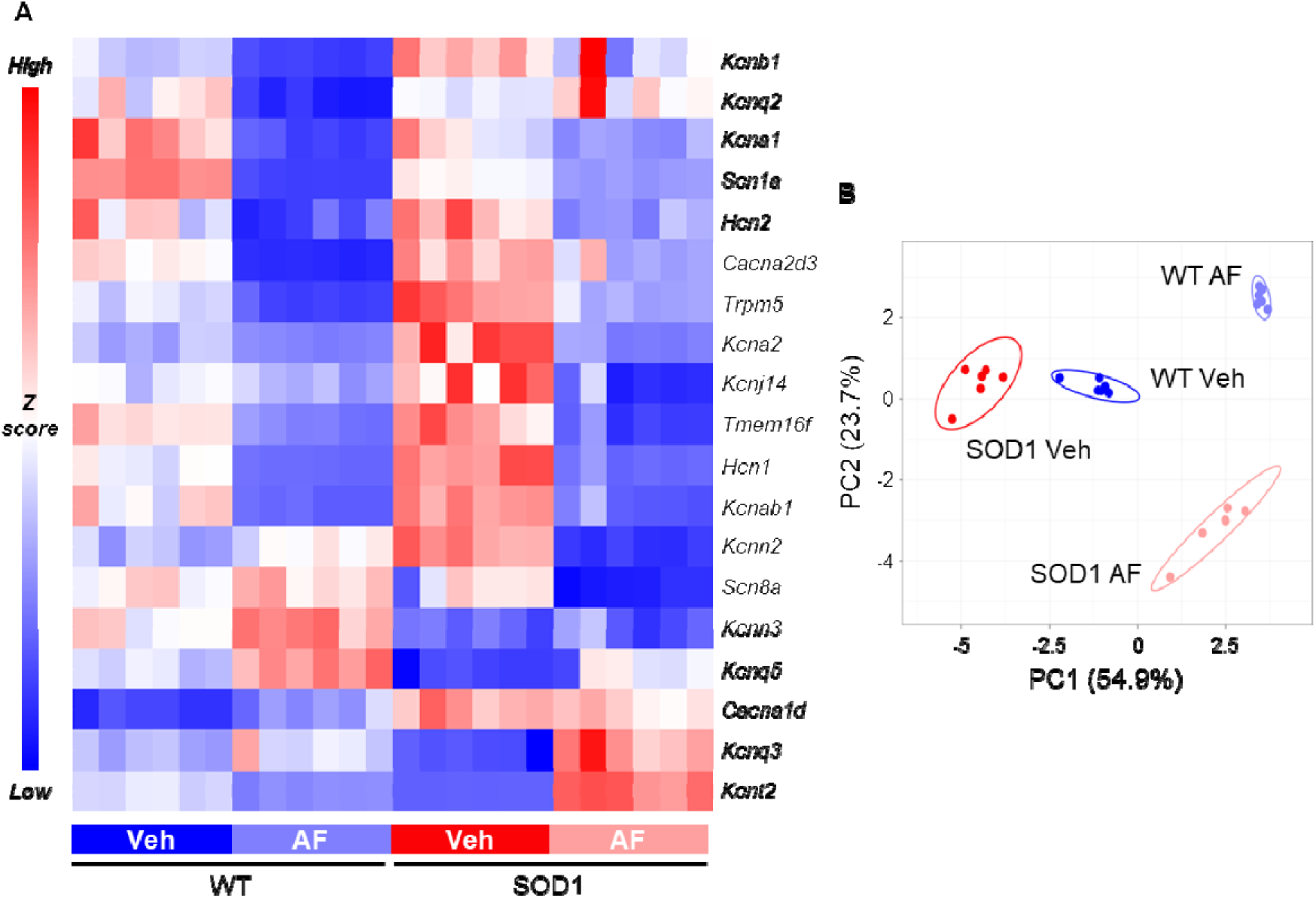
Punctual pharmacological activation of β2/β3 receptors modulates ion channel transcription in motoneurons. **A)** Targeted ion-channel transcriptome in WT and SOD1 MNs reveals the broad downregulation of multiple channels upon β2/β3 stimulation (downregulated genes*: blue*; upregulated genes: *red*). N = 6 mice per experimental group. **B)** PCA plot of ion channel gene transcription reveals the separation of experimental groups according to drug treatment and genotype. N = 6 mice per experimental group.

### 6. A prolonged delivery of β2/β3 adrenergic agonists triggers homeostatic feedback loops in MNs from WT and SOD1 presymptomatic mice

After having established how a punctual activation of the β2 and β3-adrenergic receptors modulate the MN properties, we then performed electrophysiological experiments to investigate whether a prolonged treatment during 10 days elicits homeostatic response and whether this response is similar or not in MNs from WT and SOD1 mice. We found that most of the effects of β2/β3 agonists disappeared upon chronic administration (Fig. 7, all statistics in Supplemental Excel Table 1). Gin peak (Fig. 7G), Gin plateau (Fig. 7H), and Gin ramp (Fig. 7D) were not affected by the chronic drug delivery when compared with the vehicle-treated group. V-threshold was not affected either (Fig. 7B), but the drug had a small but significant effect on Vthreshold−RMP across genotypes (Fig. 7C). However, this effect was due to a slight hyperpolarization of RMP (Fig. 7A). The recruitment current did not significantly change either upon chronic drug delivery (Fig. 7E). Most surprisingly, even the F-I gain, that was the most affected in acute conditions, did not change upon chronic conditions (Fig. 7F). To ensure the reproducibility of these findings, we repeated the *in vivo* intracellular recordings in a second independent cohort in a different laboratory and the main findings could indeed be replicated: a single delivery of β2/β3 agonists increased the F-I gain but this effect was lost after a ten days-long administration (Supplemental Fig. 1D and 1H). Altogether, these findings point to a similar homeostatic regulation of the β-adrenergic neuromodulation both in MNs from WT and SOD1 mice.

**Figure 7.**
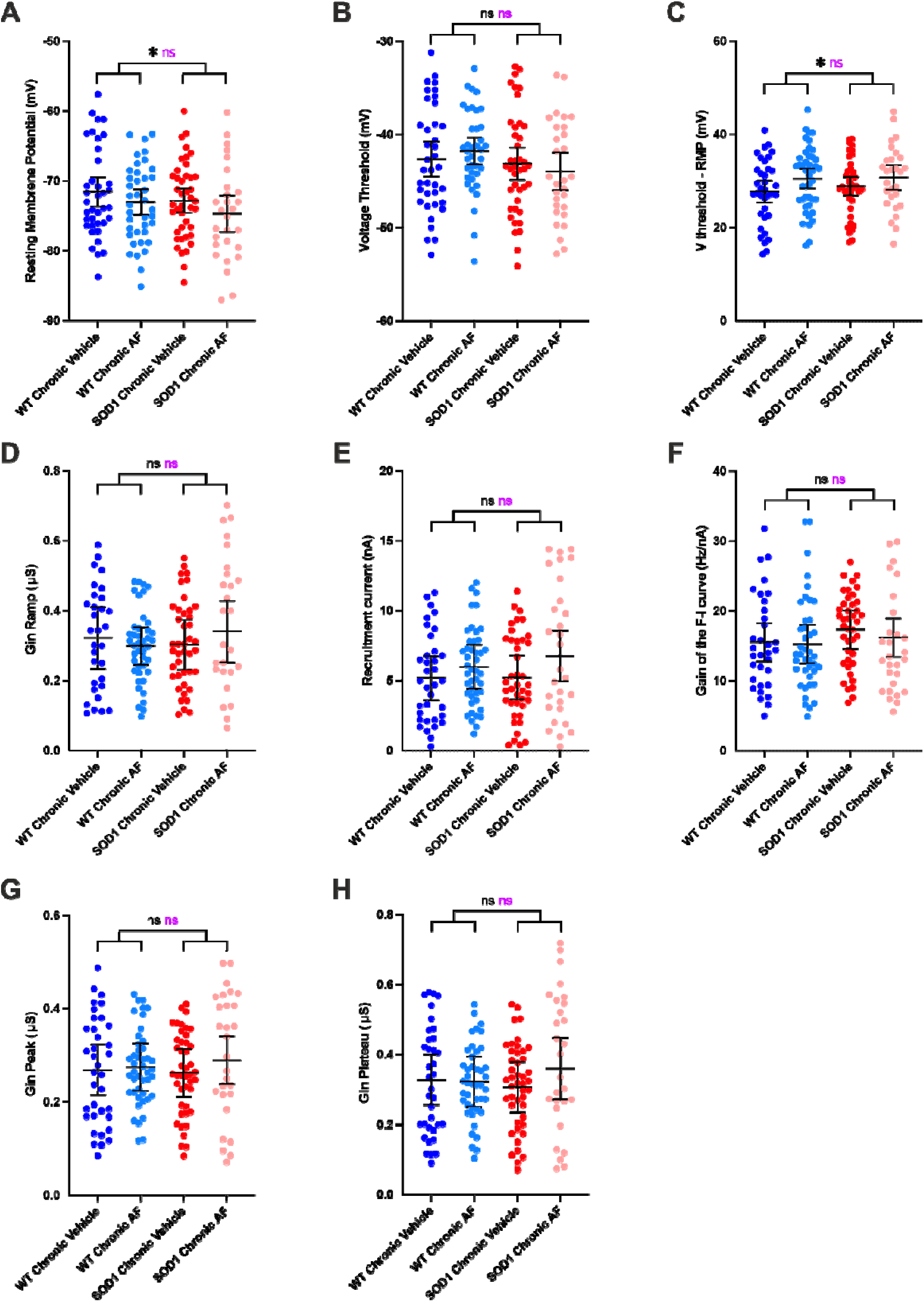
Prolonged delivery of adrenergic β2/β3 agonists does not increase the firing of motoneurons. **A-F)** Electrophysiological properties were obtained from slow ramps of current, as in Figure 4. Effect of the treatment on resting membrane potential **(A)**, voltage threshold for spiking **(B)**, Voltage threshold - resting membrane potential **(C)**, ramp input conductance **(D)**, recruitment current **(E)**, gain of the F-I relationship **(F),** in MNs from WT and SOD1 mice. **G-H)** Electrophysiological properties were obtained from a series of current pulses lasting 500ms, as in Figure 3. Effect of the chronic treatment on peak input conductance **(G)**, plateau input conductance **(H)** in MNs from WT and SOD1 mice. In all graphs, each point represents one MN and the mean ± 95% confidence intervals are shown. Significances on top bars are for treatment effects (black) and interaction effects (magenta). *Post-hoc* significances are shown for WT Chronic Vehicle *vs.* WT Chronic AF (treatment effect in WT), SOD1 Chronic Vehicle *vs.* SOD1 Chronic AF (treatment effect in SOD1) and for WT Chronic Vehicle *vs.* SOD1 Chronic Vehicle (genotype effect before treatment). N = 11 WT mice and N = 9 SOD1 mice. \**p*<0.05, ns - non significant.

In order to elucidate whether a downregulation of the GPCR themselves may contribute to this homeostatic response, we explored the effect on receptor expression either of chronic cAMP/PKA activation or of chronic β2/β3 stimulation. First, we used DREADD-Gs to stimulate cAMP/PKA in MNs (Fig. 8A-C); DREADD expression was obtained by intraspinal injection of AAV9 in SOD1/ChAT-Cre double-transgenic mice (to restrict the expression to MNs; Bączyk *et al*., 2020). Upon administration of the cognate agonist (Clozapine-N-Oxide, 10 days), we observed a substantial downregulation of *Adrb1* in MNs (by ISH; Fig. 8A, 8C) but not of the PKA-coupled *Drd5* (Fig. 8B-C). On the other hand, 10 days-long administration of β2/β3 agonists resulted in the downregulation of *Adrb1*, *Adrb2* and *Adrb3* as well as of *Drd5* genes coding for these receptors (Fig. 8D and Supplemental Excel Table 1). This reduced transcription might be the consequence of receptor desensitization and play a role in preventing the drug from increasing MN excitability. Finally, no effect was detected when evaluating the impact of chronic β2/β3 agonist treatment on MN disease markers, assessed by quantifying misfolded SOD1 levels (Fig. 8E and Supplemental Excel Table 1).

**Figure 8.**
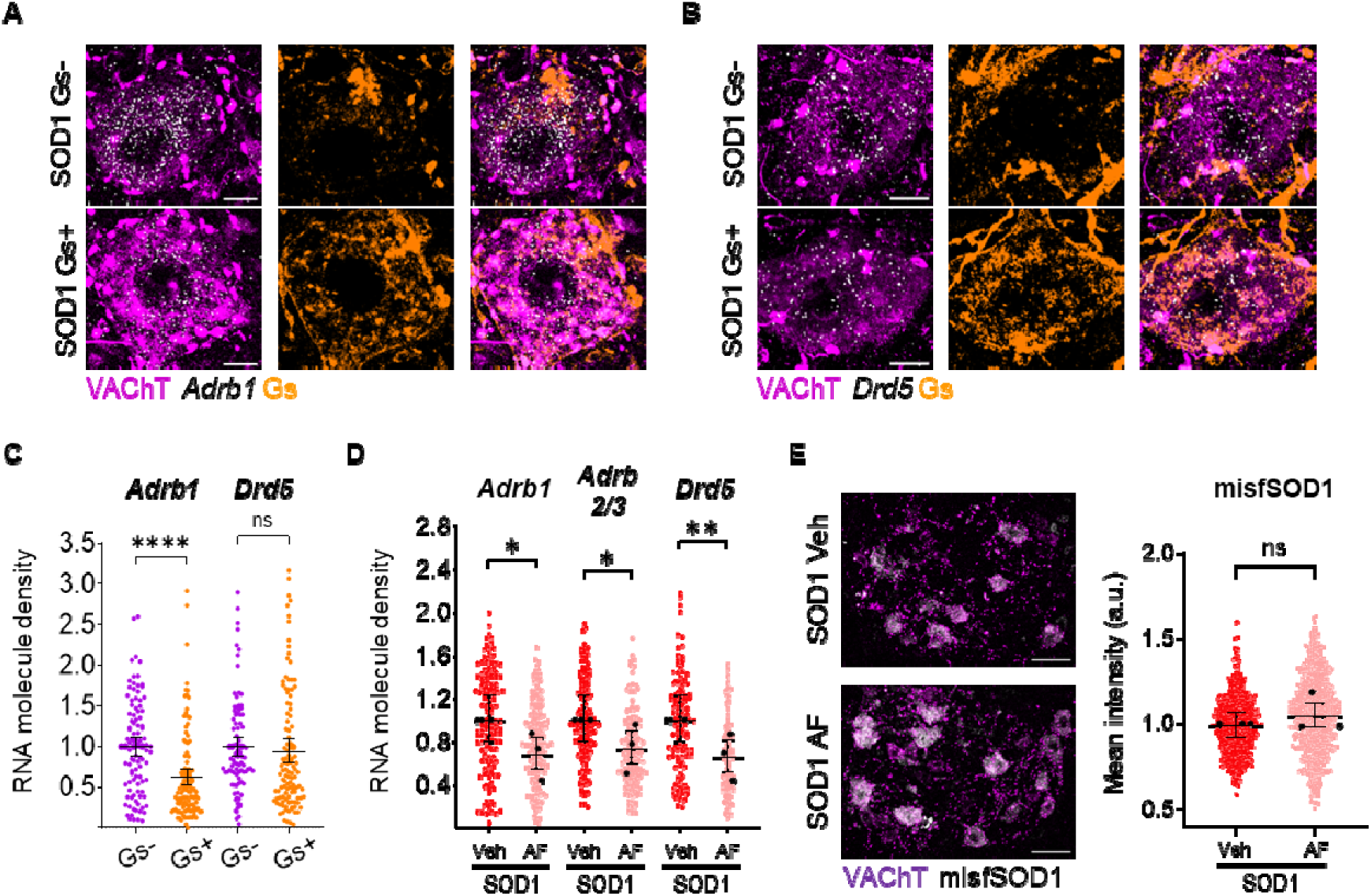
Chronic PKA engagement downregulates PKA-coupled receptors in motoneurons and does not result in amelioration of disease burden. **A-C)** DREADD(Gs)-mediated chronic chemogenetic activation of PKA pathway in presymptomatic SOD1 MNs results in the downregulation of *Adrb1* and *Drd5* mRNA. Confocal images showcase sample AAV-infected, DREADD(Gs)-expressing (Gs+) or uninfected (Gs-) MNs (from left to right: combined VAChT and receptor mRNA, DREADD(Gs), merge). Scale bar = 10µm. Individual cell expression levels are quantified as the number of mRNA molecules per µm^2^; N = 4 mice per group for *Adrb1* ISH and N = 3 mice per group for *Drd5* ISH. **D)** Expression of *Adrb1*, *Adrb2/3* and *Drd5*, revealed by single-molecule *in situ* hybridization, is downregulated by 10 days of β2/β3 agonists administration. mRNA levels are assessed as in A); N = 3 animals per experimental group. **E)** Misfolded SOD1 burden in MNs is not modified by 10 days β2/β3 agonists administration. Left panel, representative confocal images of misfolded SOD1 immunostaining in vehicle- or AF-chronically treated lumbar MNs (scale bar = 50µm). Right panel, quantification of misfolded SOD1 accumulation, assessed as mean cytoplasmic intensity per cell (arbitrary units); N = 3 mice per group. In all dotplots, small dots correspond to individual MN data points, whereas large dots represent individual animal means; 95% confidence intervals are indicated. \**p*<0.05, \*\**p*<0.01, \*\*\*\**p*<0.0001, ns - non significant.

## Discussion

This study reveals a previously overlooked β-adrenergic neuromodulation of MNs and details mechanisms of homeostatic feedback. Specifically, β2 and β3 adrenergic receptors modulate MN excitability and firing properties while being homeostatically regulated via transcriptional changes. These changes affect ion channel expression and lead to the downregulation of PKA-coupled receptors, including β2 and β3 adrenergic receptors. Notably, these homeostatic mechanisms function similarly in WT and presymptomatic adult SOD1 mice, indicating they remain intact at this disease stage.

### An overlooked **β**-adrenergic modulation of spinal MN excitability

Noradrenergic (NA) descending fibers originating from the locus coeruleus and subcoeruleus are long known to innervate the spinal cord, where they were shown to play a role in initiating the locomotion (Jordan *et al*., 2008). Noradrenaline differentially modulates the activity of spinal interneurons involved in locomotion (Bras *et al*., 1990; Hammar *et al*., 2004). Descending noradrenergic fibers also directly contact spinal MNs. Each spinal MN was found to receive more than one thousand NA contacts widely distributed throughout the dendritic arborization (Montague *et al*., 2013; Maratta *et al*., 2015). While noradrenergic neuromodulation of MN excitability through α-adrenergic receptors has been documented, neuromodulation through β-adrenergic receptors has so far received little attention. Actually, it was initially reported that MNs are endowed with α-adrenergic receptors but are deprived of β-adrenergic receptors (Nicholas *et al*., 1996). Many works reported multiple effects of noradrenaline on MN excitability (e.g., membrane depolarization, increased input resistance, hyperpolarization of the voltage threshold for spiking, increase of the F-I gain). These effects, may differ from one motor pool to another depending on its location in the brainstem or the spinal cord, but they were most of the time ascribed to α1 adrenergic receptors, sometime to α2 receptors, but never to β-receptors (see Rekling *et al*., 2000, for a review). However a more recent investigation demonstrated the presence of β1 adrenergic receptors in lumbar MNs of neonate rats and showed that their pharmacological activation decreased the F-I gain (Tartas *et al*., 2010). To our knowledge, the present study is the only one that investigated the modulation of MN excitability by β2/β3-receptors. We demonstrated that FDA-approved β2/β3 adrenergic receptors agonists increase the excitability of MNs both in WT and in presymptomatic SOD1 adult mice, primarily through an increase of the F-I gain and, secondary, through a decrease of the stationary input conductance (Gin ramp, Gin plateau) in SOD1 mice and a hyperpolarization of the voltage threshold for spiking in WT mice.

### Neuromodulation of mAHP through **β**2/**β**3 adrenergic receptors

It was established that the medium afterhyperpolarization (mAHP), which follows each action potential, is a potent controller of the MN firing, in particular of the gain of the F-I relationship in primary range (Kernell, 1999; Manuel *et al*., 2006). Indeed, in simple MN models, the F-I gain in primary range depends on the inverse of the product between AHP conductance, AHP decay time and AHP driving force (Meunier and Borejsza, 2005; Manuel *et al*., 2006). The smaller and faster is the mAHP conductance, the higher is the F-I gain in the primary range. We found here that β2/β3 adrenergic receptors agonists shortened the AHP half decay time indicating a faster kinetics of the mAHP conductance that accounts for, at least partly, the increase of the F-I gain. Furthermore, β2/β3 agonists reduce the voltage threshold for spiking in MNs from WT mice decreasing the AHP driving force that contributes to increase the F-I gain in these MNs. A secondary determinant of the frequency-current gain is the plateau input conductance (Meunier and Borejsza, 2005) that was found to be reduced by β2/β3 agonists in MNs from SOD1 mice. In total the reduced AHP half decay time, voltage threshold for spiking and Gin plateau may all contribute to the increased frequency-current gain.

The mAHP current in rodent MNs is mainly caused by the SK2 and SK3 calcium-dependent potassium channels, which were found to cluster on soma and proximal dendrites of α-MNs at the level of C-boutons (Deardorff *et al*., 2013). Indeed, blockade of SK channels with apamin increases the gain of the F-I relationship in spinal MNs (Zhang and Krnjević, 1987). How do β2/β3 receptors act on the SK channels? SK channels display constitutive trafficking, and PKA activation was shown to decrease the surface expression of SK channels through three phosphorylation sites (Ser568, Ser569 and Ser570 at the carboxyl-terminal region) (Ren *et al*., 2006). In the present work, we showed that an intracellular injection of cAMP-SP (a cAMP agonist) also increases the F-I gain relationship, similarly to the systemic delivery of β2/β3 agonists. Furthermore, we also showed that the pharmacological stimulation of the β2/β3 adrenergic Gs-coupled receptors effectively engages the cAMP/PKA pathway. We then speculate that the pharmacological stimulation of β2/β3 adrenergic receptors indirectly removes SK channels from the plasma membrane of MNs through the activation of the cAMP/PKA pathway. A similar scenario was previously demonstrated at excitatory synapses from lateral amygdala pyramidal neurons (Faber *et al*., 2008). This would result in a decrease of the SK conductance, entailing an increase of the F-I gain in the primary range, as observed in our experiments.

### Chronic **β**-adrenergic stimulation results in homeostatic mechanisms that cancel physiological neuromodulation

Prolonged β2/β3 agonists treatment of either WT or SOD1 animals results, in our experiments, in the loss of their neuromodulatory effects on MN electrophysiological properties. This homeostatic response may be caused by several mechanisms at transcriptional, post-transcriptional and post-translational levels. 1) We show that the β2/β3 mRNA is downregulated upon chronic pharmacological stimulation of the β2/β3 receptors establishing a negative feedback loop. Notably, heterologous downregulation of other Gs-coupled GPCRs is observed upon β2/β3 agonists treatment (or chemogenetic stimulation of Gs signaling). Our findings are in agreement with the downregulation of adrenergic receptors mRNA in multiple organs upon chronic adrenergic stimulation (Mak *et al*., 1995; Lefkowitz *et al*., 2000; Milano *et al*., 2018) through mechanisms involving transcriptional regulation and mRNA stability (Danner *et al*., 1998), as well as with extensive heteronomous downregulation of Gs-coupled GPCRs. 2) A single injection of β2/β3 agonists rapidly and substantially modulates the expression of many channel-encoding genes in MNs, often in directions that oppose the effects of β2/β3 agonists on increasing MN excitability and firing. In particular, *Scn1a* (Na_V_1.1, sodium current) is strongly downregulated while *kcnq3* (K_V_7.3) and *kcnq5* (K_v_7.5), which mediate the M-current, are upregulated (although not *kcnq2*) in both WT and SOD1 MNs contributing to reduce MN excitability (Sharples *et al*., 2023; Schroeder *et al*., 2000). On the same line β2/β3 agonists downregulate*Tmem16f* counteracting the recruitment current decrease in both WT and SOD1 MNs, (Soulard *et al*., 2020). Furthermore, while β2/β3 agonists decrease the AHP decay time, *Kcnn2* (SK2) is significantly upregulated in WT MNs (although not in SOD1 MNs), resulting in larger SK2 conductance that may increase the AHP decay time and counteract the F-I gain increase. While these transcriptional changes may only partially translate into protein-level changes within the time frame of the acute experiments (up to 3 hours), they would most likely contribute to the homeostatic feedback after the chronic pharmacological treatment (10 days). 3) Post-translational mechanisms may also contribute to stabilizing β-adrenergic signaling against persistent stimulation. Upon activation, phosphorylation of the cytoplasmic tail of adrenergic receptors results in their interaction with β-arrestin adaptors that mediate their endosomal internalization and thereby signal desensitization (Hausdorff *et al*., 1991; Tran *et al*., 2004; Wachter and Gilbert, 2012) in a timeframe of minutes to hours. Moreover, persistent stimulation leads to the phosphorylation of the receptors by β-adrenergic receptor kinases (BARKs) triggering a switch in the affinity from Gs to Gi and the reversal of the Adenylyl Cyclase activation (Wang *et al*., 2017). These post-translational homeostatic mechanisms extend to other convergent receptors through heterologous desensitization mechanisms involving PKA phosphorylation (Benovic *et al*., 1985; Gainetdinov *et al*., 2004). To sum up, our findings show that, both healthy and ALS MNs display substantial homeostatic loops regulating PKA signaling and excitability responses to GPCR stimulation. Furthermore, the homeostatic regulation of the β-adrenergic neuromodulation upon prolonged agonist administration is the same in MNs from SOD1 and WT mice indicating that MNs are still displaying a proper gain in the feedback loops and a meaningful homeostatic regulation of excitability at a presymptomatic disease stage.

## Conclusions

Our findings demonstrate that MNs are responsive to neuromodulatory signals not only from α- but also from β-adrenergic receptors, hinting at an integration of these two distinct adrenergic signaling cascades with each other and possibly with other neuromodulators. We demonstrate that homeostatic processes, at receptor and ion-channel levels, are set in motion by β-adrenergic stimulation aimed at restoring an excitability set-point both in normal and disease conditions.

## Acknowledgments

The authors acknowledge the animal facility of BioMedTech *Facilities* at Université Paris Cité (INSERM US36 / CNRS UAR2009) for its support and expertise. We thank the Thierry Latran Foundation (“TRiALS” project), the National Institutes of Health, National Institute of Neurological Disorders and Stroke (R01NS110953, R01NS115900, R01NS112304), the join Deutsche Forschungsgemeinschaft/Agence Nationale de la Recherche “SynaptALS” project (ANR-20-CE92-0029-01/ DFG 446067541) and Polish National Science Centre (OPUS 2019/35/B/NZ4/02058) for their financial support.

## Authors contributions

Conceptualization, M.B., F.R., D.Z., Methodology, S.A., M.B., G.C., S.D., F.R., Software, S.A., S.D., Formal Analysis, S.D., Investigation, S.A, N.D., M.B., A.T., F.o.H, H.Z., G.C., K.G., S.S.K., Data Curation, G.C., S.D., Writing-original draft, S.A., M.B., G.C., S.D., F.R., D.Z., Writing - Review & Editing, A.L., S.A., M.B., G.C., S.D., F.R., D.Z., Visualization, S.A., G.C., Supervision, M.B., F.R., D.Z., Funding Acquisition, A.L., M.B., F.R., D.Z.

## Methods

### Animals

All experiments were performed on the transgenic mouse line B6SJL-Tg(SOD1*G93A) 1Gur/J pursued via The Jackson Laboratory (#002726, Gurney *et al*., 1994). SOD1*G93A hemizygotes (here referred to as SOD1) develop a phenotype resembling human ALS as a consequence of the high number of transgene copies. Only males were used in Ulm (histology and transcriptomics experiments) in accordance with literature characterizing disease staging in this mouse strain (Gurney *et al*., 1994; Saxena *et al*., 2013; Ouali Alami *et al*., 2018; Bączyk *et al*., 2020), animals aged P40-P45 were deemed presymptomatic. Animal experiments from Ulm were conducted according to institutional guidelines (Tierforschungszentrum, Universität Ulm, Germany) under the approval of the Regierungspräsidium Tübingen with license nr. 1390 (untreated) and 1440 (drug-treated animals).

Animal experiments from Paris were approved by the Paris Descartes University ethics committee (CEEA34) and authorized by the French ministry for higher education and research (authorization number APAFIS#16338-2018052100307589). Animal experiments from Poznań were approved by the local ethical committee (16/2021). For electrophysiological experiments in Paris and in Poznań, both male and female mice were recorded between the ages of P45-P59.

Mice were grouped ≤5 with access to food and water *ad libitum* and housed in M2 long cages in an open shelving system (Ulm, Poznań) or disposable and ventilated cages (Paris) under a 12h/12h light/dark cycle with 40÷60% relative humidity. Upon routine tests assessing motor impairment, individuals reaching end-stage (on average >P110-P120) were euthanized complying with the aforementioned directives. In Paris, SOD1 mice were euthanized no later than P90.

For electrophysiological experiments B6SJL-Tg(SOD1*G93A)1Gur/J males were crossed with B6SJL/J females, and offspring carriers of the mutated SOD1 gene were selected for experimental groups (SOD1), with non-transgenic littermates serving as wild type (WT) control animals. For chemogenetics experiments (Regierungspräsidium Tübingen, license nr. 1404), females from the B6;129S6-ChAT tm2(cre)Lowl /J line (The Jackson Laboratory, stock #006410; Rossi *et al*., 2011) were crossed with SOD1 males; only F1 SOD1;ChAT-Cre males were used for intraspinal injections of AAV9 viruses enabling Cre-dependent expression of the Gs-DREADD construct in ChAT+ cells (viruses were produced as in Commisso *et al*., 2018).

### Surgical procedures for electrophysiological experiments

In Paris and Poznań, electrophysiological experiments were carried out with the same protocol. Fifteen minutes before anesthesia, atropine (0.20mg/kg; Aguettant or Polfa) and methylprednisolone (0.05mg; Solu-Medrol; Pfizer) were given subcutaneously to prevent salivation and edema, respectively. Anesthesia was induced with an intraperitoneal (i.p.) injection of Fentanyl 0.025mg/kg, Midazolam 7.5mg/kg and Medetomidine 0.5mg/kg. The heart rate was monitored with an EKG, and the central temperature was kept around 37°C using an infrared heating lamp and an electric blanket. Then, the mouse was artificially ventilated with pure oxygen (SAR-1000 ventilator; CWE) through a cannula inserted in the trachea. The ventilator settings were adjusted to maintain the end-tidal CO_2_ level at ∼4% (Micro-Capstar; CWE). Two catheters were introduced in the external jugular veins. The first one was used to deliver supplemental doses of anesthesia whenever necessary (usually every 20–30 min) by i.v. injection (10% of the dose used for anesthesia induction). The adequacy of anesthesia was assessed by lack of noxious reflexes and stability of the heart rate (usually 400–500 bpm) and end-tidal pCO_2_. The other catheter was used to slowly inject (50μL/h) a 4% glucose solution containing NaHCO_3_ (1%) and gelatin (15%; Voluven; Fresenius Kabi or Tetraspan; Braun) to maintain the physiological parameters and to inject the tested treatment. The sciatic nerve was dissected and mounted on a bipolar electrode for stimulation. The vertebral column was immobilized with two pairs of horizontal bars (Cunningham Spinal Adaptor; Stoelting) applied on the Th12 and L2 vertebral bodies, and the L3–L4 spinal segments were exposed by a laminectomy at the Th13–L1 level. The exposed tissues from the hindlimb and spinal cord were covered with pools of mineral oil. When the surgery was completed, the animal was paralyzed with vecuronium or pancuronium with a bolus of 0.5mg/kg as needed. Additional doses of anesthetic were then provided at the same frequency as before the paralysis, and adequacy of anesthesia was assessed by stability of the heart rate and pCO_2_. At the end of the experiment, animals were euthanized with a lethal i.p. injection of pentobarbital (Exagon or Morbital; 80mg).

### Stimulation and intracellular recordings

Intracellular recordings of MNs were performed with micropipettes (tip diameter, 1.0– 1.5μm) filled with 2M K-acetate (resistance □25MΩ). Recordings were made using an Axoclamp 2B (Paris) or Axoclamp 900A (Poznań) amplifier (Molecular Devices) connected to a Power1401 interface (sampling rate 20kHz) and using Spike2 software (CED). After impalement, identification of MNs relied on the observation of antidromic action potentials in response to the electrical stimulation of the sciatic nerve. All MNs retained for analysis had a resting membrane potential more hyperpolarized than −50mV and an overshooting action potential >65mV. For more details on the analysis, see Manuel *et al*. (2009).

Briefly, the input conductance was determined from the I-V relationship, which was plotted from the peak (Gin peak) and plateau (Gin plateau) responses to a series of small-amplitude square current pulses (−2 to +2nA, 500ms). Discharge properties were investigated using slow triangular ramps of currents (0.5 - 2nA/s). The discharge frequency-current relationship (F-I) was determined by plotting the instantaneous firing frequency against the current intensity. The input conductance was also determined from the early linear voltage response (<2nA) to a slow ramp of current (Gin ramp). All recordings were performed in discontinuous current clamp mode (8kHz). All care was taken to compensate for the microelectrode resistance and capacitance.

### PKA activation experiments

The procedures for surgery, MN stimulation and recording were performed as described above. To measure the impact of PKA activation on MN intrinsic excitability, the recording microelectrode was filled with a mixture of 2M K-acetate, and 4mM cAMP analogue (S)-adenosine, cyclic 39,59- (hydrogenphosphorothioate) triethylammonium (cAMP-SP; Sigma). Upon penetration of a MN, series of square current pulses and slow ramp of currents were repeated in successive 3-5 min intervals for at least 15 min, until the properties of the MN stabilized. Then, the iontophoretic injection of cAMP-SP was performed through the microelectrode (500ms pulses of −2 to −4nA, repeated at 1Hz, total normalized electrical load injected 2600 ± 600nA.sec/µS. We adjusted the injected current to the MN input conductance (normalized electrical load) because we noticed that MNs with higher conductance needed more current injection to observe the same effects as in MNs with smaller conductance. Immediately after, the square current pulses and slow ramp of currents were repeated with the same parameters as before the activation of the cAMP/PKA pathway. Data were excluded when the resting membrane potential changed by more than ±5mV or the bridge balance changed by more than ± 20%, to ensure that the changes in excitability are not due to an insufficient stability of the MN or the microelectrode.

### Surgical procedures for intraspinal AAV injections

Intraspinal surgery and AAV injection procedures were performed as in Bączyk *et al*. (2020). In short, P18 to P21 mice first received a subcutaneous dose of buprenorphine (0.1mg/kg) and meloxicam (1mg/kg) and from then onwards they were kept anesthetized onto the stereotaxic frame via continuous administration of 2% isoflurane in O_2_ at 0.8L/min O_2_ flow rate. After making a small incision on the skin, the subcutaneous fascia was removed and the paraspinal muscles were gently pushed sideways and blunt-dissected. The underlying spinal cord was exposed by conducting laminectomy on T12 and removing the vertebral bone flap. Using a Picospritzer microfluidic device, 1μL of a 1:1 mixture of virus suspension and 1% Fast Green dye in PBS++ was injected with a pulled glass capillary at the following stereotaxic coordinates: y=+0.25mm with respect to the dorsal artery; 0.4mm depth below the dorsal surface of the spinal cord. Before withdrawing it, the capillary was left in place for 10 minutes to allow the viral suspension to diffuse locally and to avoid viral backflow. Over the next 3 days, mice were treated with meloxicam (daily, 1mg/kg) and buprenorphine (twice a day, 0.1mg/kg) and monitored for post-operative complications.

### Drug treatments

The Adrb ligands selected for acute or chronic (10 days-long) intraperitoneal (i.p.) treatment were amibegron (A; Cayman Chemicals 11954), an Adrb3 agonist, and formoterol fumarate (F; LKT Laboratories F5868), an Adrb2 agonist. All drugs were diluted in a mixture of 5% dimethyl sulfoxide (DMSO) 5% Tween-80 5% polyethylenglycole-400 (Roth) in saline to achieve the designated daily dosage (AF, 10mg/kg A and 0.3mg/kg F). Intracardiac perfusions took place 3h after the latest injection.

For electrophysiological experiments, A and F stock solutions were separately diluted in DMSO (Sigma) and stored at −20°C until the day of the experiment. The day of the experiment, stock solutions were diluted in saline (B.Braun) and mixed together to obtain the desired dose (A: 3mg/kg and F: 0.3mg/kg). The AF cocktail was injected i.v. after at least one MN was recorded. Just after the injection of AF, an increased heart rate was clearly observable on the EKG and started to decrease after 3h. Because β-adrenergic receptors are expressed in the heart and are known to increase the heart rate (Lohse *et al*., 2003), we used the heart rate as an indicator of an ongoing activation of β-adrenergic receptors. Therefore, MNs were recorded and kept for analysis for up to 3h after the injection of AF.

For chemogenetics experiments, mice injected with Gs-DREADD-expressing AAV9 were administered the Gs-DREADD agonist clozapine N-oxide (CNO) between P35 and P45 i.p. route twice per day at 5mg/kg, with an 8h interval between consecutive treatments. Drugs were solubilized in DMSO and diluted in saline (2.5% DMSO in the final mixture). Intracardiac perfusions took place 3h after the last injection.

### Laser capture microdissection and RT-qPCR

Mice received terminal anesthesia by i.p. administration of ketamine (100mg/kg; WDT) and xylazine (16mg/kg; Bayer) and were perfused with RNAse-free ice-cold PBS for 2 min at 7mL/min flow rate. The lumbar portion of the spinal cord was quickly dissected out, OCT embedded (Sakura) and stored at −80°C. 12μm-thick cryosections were converted onto RNAse-free polyethylene terephthalate membrane (PET) slides (Zeiss 415190-9051-000) once these had been coated with poly-L-Lysine (Sigma Aldrich).

Firstly, a fixation step was performed in −20°C-cold 70% ethanol in RNAse-free H_2_O (DEPC ddH_2_O: 1% diethylpyrocarbonate (Roth) in ddH_2_O). Next, the slides were stained for 1 min in 4°C-cold 1% cresyl violet (WALDECK) in 50% ethanol-DEPC ddH_2_O and finally washed for 1 min in 4°C-cold 70% and then 100% ethanol-DEPC ddH_2_O. 30 MNs per group were microdissected into 500µL adhesive caps (Zeiss) with a laser microdissection system (Palm MicroBeam, Zeiss). MNs were lysed by pipetting 21μL of a mixture comprising 30μL 10X RT buffer, 3μL RNAse OUT (Invitrogen™ SuperScript™ III Reverse Transcriptase kit, 11904018) and 0.3μL 1% NP-40 in ddH_2_O directly onto the adhesive cap. After vortexing upside-down for 30s and incubating for 20 min at 42°C, samples were frozen at −80°C overnight.

Reverse transcription and RNA digestion were carried out with the cDNA synthesis kit as per manufacturer’s instructions. RNA content was determined via qPCR with a Roche LC480 cycler using 2μL cDNA. PCR cycles were set as follows: 2 min at 50°C, 10□min at 95°C (initial denaturation); 50 cycles comprising 15s at 95°C (denaturation) and 1 min at 60°C for annealing and elongation. The relative quantification of the transcripts of interest relied on the ΔCt method upon normalization against *Gapdh* housekeeping gene; the sequences of the primers used in this study, designed with NCBI Primer designing tool (Primer-BLAST, Ye *et al*., 2012) are reported in Supplemental Table 1.

### Histology

Terminally anesthetized P45 animals were perfused first with 50mL ice-cold PBS (prepared in house) and 50mL 4% paraformaldehyde (PFA; Sigma-Aldrich) in PBS at an approx. 7mL/min flow speed. Overnight fixation in 4% PFA in PBS preceded incubation in 30% sucrose (Roth) in PBS and OCT embedding. Samples were equilibrated overnight at −20°C before being sequentially cut into 15μm-thick sections.

*In situ* hybridization (ISH) experiments were carried out with the RNAscope Fluorescent multiplex reagent kit v1 (Advanced Cell Diagnostics, ACD; Wang *et al*., 2012) and the following ACD probes: Mmp9-C1 (315941), Adrb1-C3 (449761), Adora2b-C3 (445281), Drd5-C3 (494411), Adrb2-C3 (449771), Adrb3-O1-C3 (502581). The manufacturer protocol for fixed frozen tissue was followed with minor deviations (for detailed procedures, refer to olde Heuvel *et al*., 2019).

Subsequently, sections were co-immunostained to identify α-MNs; to prevent ISH signal disruption, the slides were always incubated in a humid chamber and protected from light. In short, sections were washed twice in 1X Wash Buffer for 10 min at room temperature (RT), incubated for 1h at RT in blocking buffer (BB: 10% Bovine Serum Albumin 0.3% Triton™X-100 (Sigma-Aldrich) in 1X PBS) and finally with primary antibodies overnight at 4°C. In this manuscript primary antibodies (anti-VAChT, Synaptic Systems, 139105; anti-mCherry, NanoTag Biotechnologies, N0404-AT565) were pre-diluted 1:250 in BB. Three 30 min washes in 0.1% Triton™X-100 in 1X PBS (PBS-T) at RT preceded incubation with pre-diluted secondary antibodies (1:250 in BB) for 2h at RT. All secondary antibodies (Sigma, SAB4600468 used for co-immunostaining were applied at 1:250 dilution. For free-floating immunostaining experiments, sections were incubated in BB for 2h and with primary antibodies (anti-misfSOD1 1:1000, Médimabs MM-0070; anti-VAChT 1:500, Synaptic Systems, 139105; anti-phospho-PKA substrate 1:100, CST 9621) for 48h; secondary antibodies (Invitrogen, A21202; Biotium, 20171; Invitrogen, A32790) were applied at 1:500 dilution.

### Imaging

All experiments comprising an ISH step were imaged using a confocal laser scanning microscope (Carl Zeiss LSM710) equipped with a Plan-Apochromat 63X oil objective (NA 1.40). Images were acquired with a 1024×1024 frame size at 12-*bit* depth (0.132μm/px resolution in *xy*; 0.57μm voxel depth) preferentially at the surface of the section where probe penetration was highest and thus consistent between groups. The same image acquisition settings were employed to image misfSOD1 accumulation and phospho-PKA epitopes, but with a 20X objective (NA 0.8). Only ventrolateral MNs were investigated in this study.

### Image processing and quantification

Maximum intensity projections of confocal *z*-stacks and in general all downstream processing and image analysis steps were instead performed in *Fiji* (Schindelin *et al*., 2012). *ImageJ* custom macros enabled semi-automated identification, drawing and ID assignment of MN *somata* according to VAChT signal. Regions of interest were then extracted from ISH channels (upon a background subtraction step, *radius* 5) and their size was measured retaining information on the receptor probe, the experimental group and the MN ID in their name tag. A similar approach was adopted to measure cytoplasmic PKA phospho-epitopes and whole-cell misfSOD1 accumulation as mean fluorescence intensity per MN cross section area. ISH signal was quantified as spot density per μm^2^ by means of custom Jython and *ImageJ* macro scripts exploiting the difference of Gaussian detector shipped with Trackmate plugin (Tinevez *et al*., 2017); peak size and intensity threshold were set upon visual inspection of the experimental group given the slight variability and efficiency of probe penetration in different batches.

### Data analysis and statistics

All RT-qPCR data were first normalized against *Gapdh* mRNA levels and, subsequently, against the control group (WT group for the LCM-based GPCR screening; WT vehicle for acute drug treatment experiments; SOD1 vehicle for chronic administration experiments). To test whether MN mRNA expression differed between WT and SOD1 mice, a *t*-test for each gene was computed. Heatmaps of LCM-based data were generated with the *ggplot2* R package.

All ISH and misfSOD1 immunohistochemistry data were first normalized against the control group for a meaningful comparison of the experimental sets; a quality-control step was further included to determine MN cross-section areas and assess whether the sampling was uniform or otherwise introducing unintended bias. Upon visual inspection of receptorome scatterplots (*Mmp9 vs.* receptor mRNA, *data not shown*), for all screenings a threshold of 0.3 spots per μm^2^ was chosen in order to subdivide MNs into *Mmp9*+ and *Mmp9*- categories. *Adrb2* and *Adrb3* data were handled with the *tidyverse* R package. The heatmap was computed after gene-wise z score data normalization in R.

For GPCR ISH screening, to test whether MN mRNA expression differed between WT and SOD1 mice and whether this effect was different between *Mmp9*- and *Mmp9*+ MNs, a generalized linear mixed model was calculated for each gene. Fixed effects for genotype (WT or SOD1) and motoneuron type (*Mmp9*- or *Mmp9*+), their interaction effect, a random per-animal offset, and a full-factorial dispersion model were included. A gaussian error distribution with an identity-link was used for *Adrb1* and *Adrb2*, whereas for *Adrb3* a Gamma distribution with a log-link was necessary to meet model assumptions.

For immunohistology of RXXpS/T, to test whether the acute treatment (vehicle or agonist) affected PKA-phosphorylation in MNs and whether this effect differed between WT and SOD1 mice, a linear mixed model with genotype, treatment and their interaction as fixed effects and a per-animal random offset was calculated.

For immediate early genes and ion channels RT-qPCR, to test if the acute treatment (vehicle or agonist) affected MN mRNA expression and whether this effect differed between WT and SOD1 mice, a linear model with genotype and treatment as well as their interaction were calculated for each gene. The heatmap was computed after gene-wise z score data normalization in R, whereas the PCA plot was generated with Clustvis software (Metsalu and Vilo, 2015).

For GPCR ISH upon chemogenetic treatment, to test whether the chronic DREADD(Gs) activation affected MN transcription differed in infected (Gs+) or in non-infected (Gs-) MNs, a generalized linear mixed model with MN type (Gs+ or Gs-) as a fixed and dispersion effect, a per-animal random offset, and a Gamma error-distribution with a log-link function were calculated for each gene.

For GPCR ISH upon adrenergic agonist treatment, to test whether the chronic treatment (vehicle or agonist) affected MN transcription, a generalized linear mixed model with treatment as a fixed and dispersion effect, a per-animal random offset, and a log-link function were calculated for each gene. For *Adora2b*, a Gamma error distribution was used.

For misfSOD1 immunohistology, to test whether the chronic treatment affected misfSOD1 levels, a generalized linear mixed model with treatment as a fixed and dispersion effect, a per-animal random offset, and a log-link function was calculated.

For electrophysiology, to test if the treatment affected intrinsic neuronal properties and whether this effect differed between WT and SOD1 mice, we calculated linear mixed models for each intrinsic parameter. Fixed effects for treatment (acute: before or after; chronic: vehicle or agonist) and genotype (WT or SOD1) as well as their interaction effect were included. To account for between animal variance, a random per-animal offset was introduced. Backwards elimination was performed on the random effect, *i.e.*, the random offset was only included if it significantly contributed to goodness-of-fit of the model.

Data obtained from the same MNs recorded before and after the iontophoretic injection of cAMP-SP were compared using a paired *t*-test except for maximum firing frequency, for which a Wilcoxon paired test was used.

Model assumptions were confirmed both by visually inspecting the quantile-quantile plots and histograms, and by performing distribution, dispersion, outlier, homogeneity of variance, and quantile deviation tests using the DHARMa R package (Hartig, 2022). If model assumptions were violated, following strategies were employed. Data points were identified as outliers, if the absolute value of their residual was greater than 2 standard deviations, and removed. In case of non-normality of residuals, generalized-linear (mixed) models with link function and error distributions that did not violate model assumptions were chosen. A full-factorial dispersion model was fit in case of presence of heteroscedasticity. Degrees of freedom of linear mixed models were adjusted using the Kenward-Roger method.

Linear models were fit using lm, linear mixed models using *lmer* of the *lme4* R-package (Bates *et al*., 2015), and generalized linear (mixed) models using *glmmTMB* (a Template Model Builder interface for R; Brooks *et al*., 2017). Planned *post-hoc* tests were performed to test if there was a significant treatment effect within WT or SOD1 mice. Throughout, an alpha-error of *p*<0.05 was regarded as significant.

---

**Supplemental Table 1.**
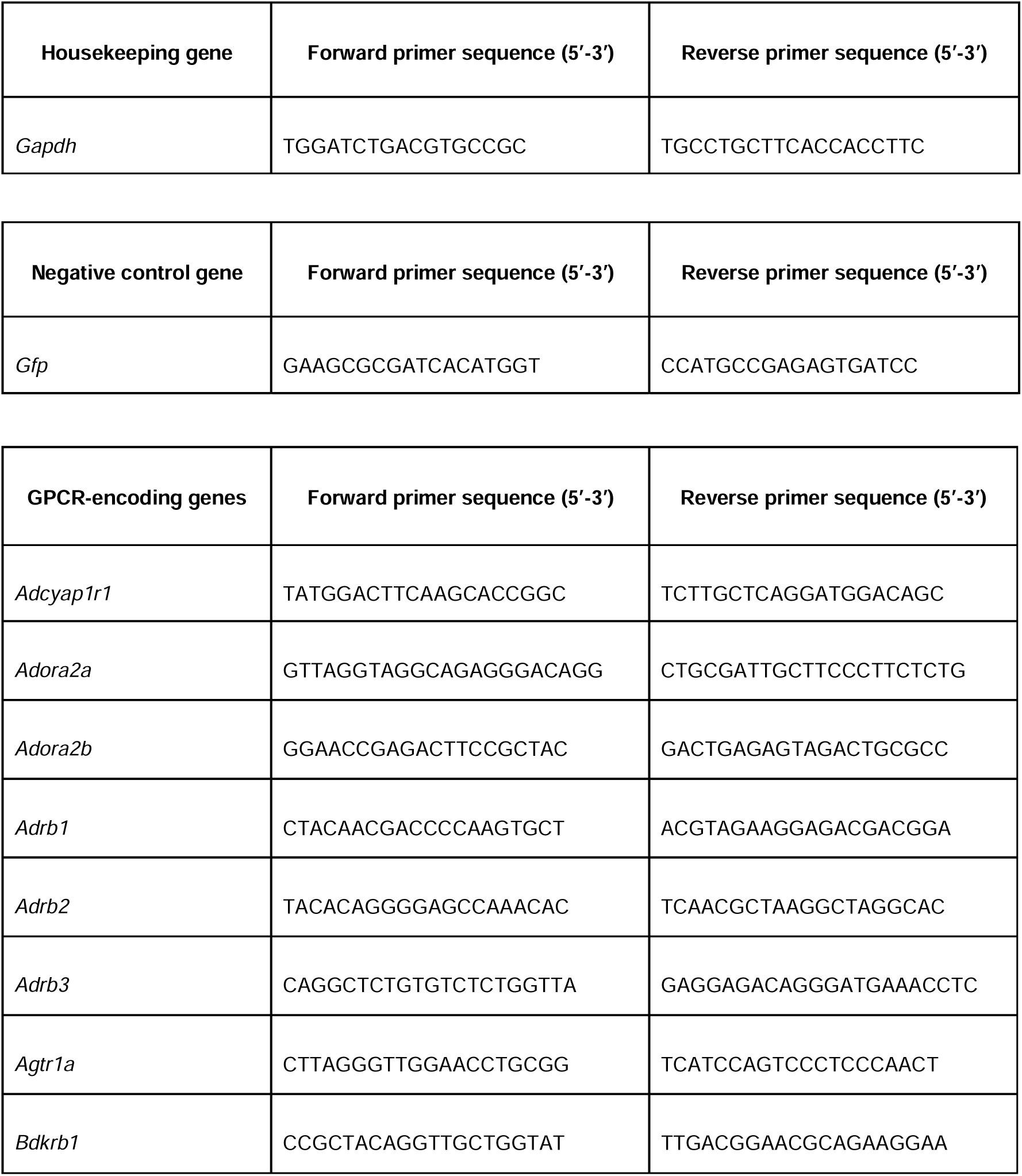
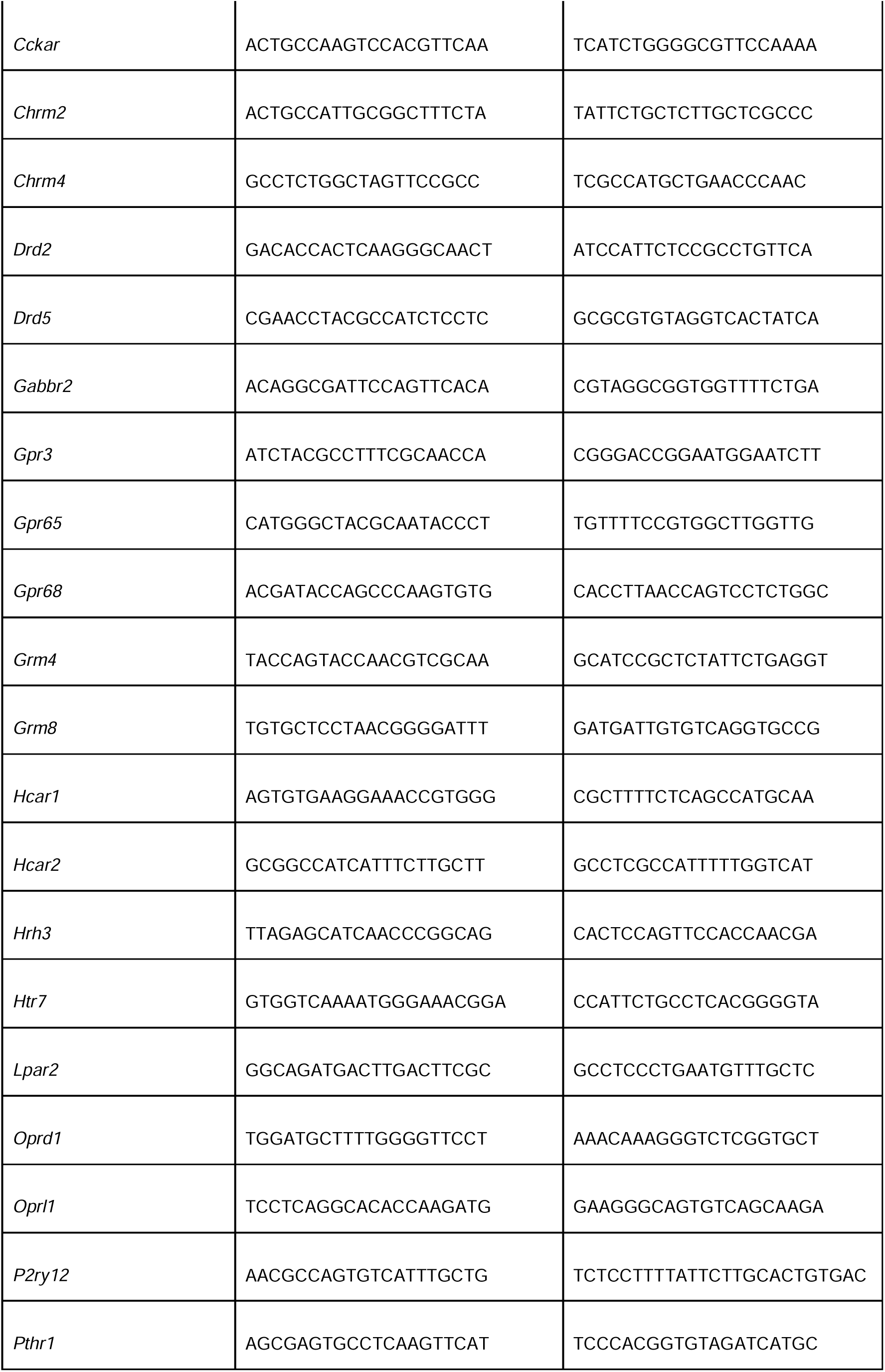
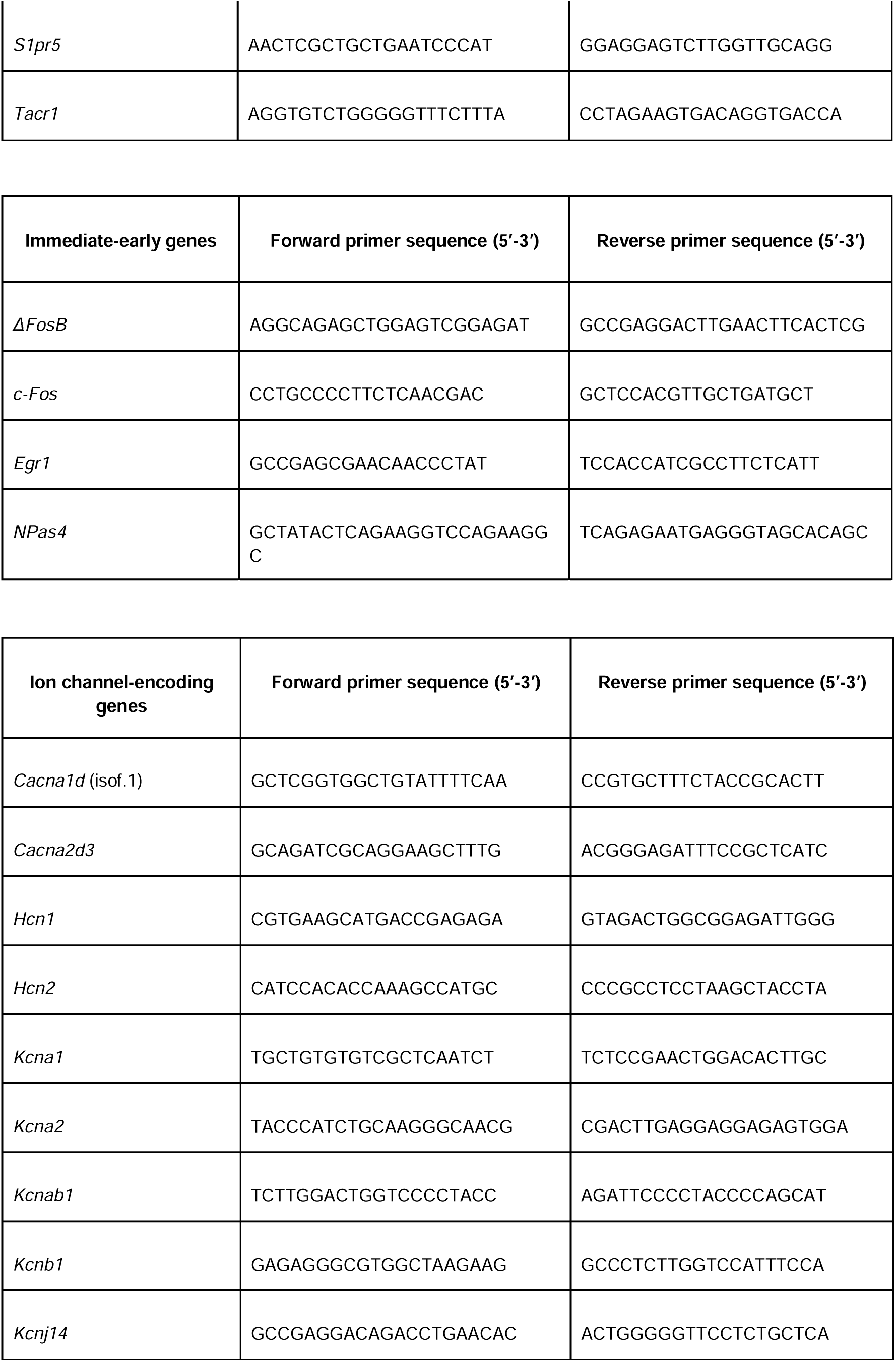
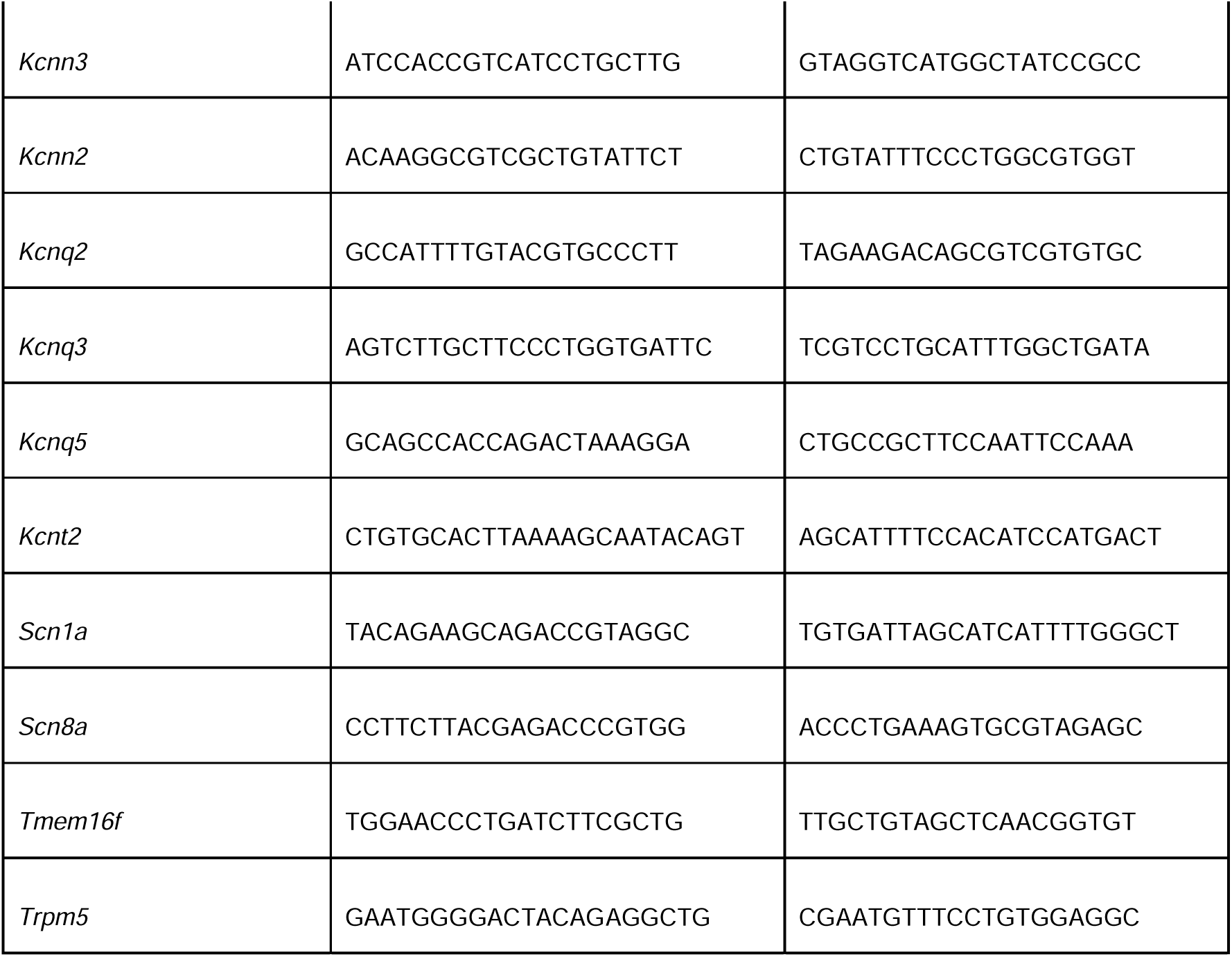
qPCR primer list.

**Supplemental Figure 1.**
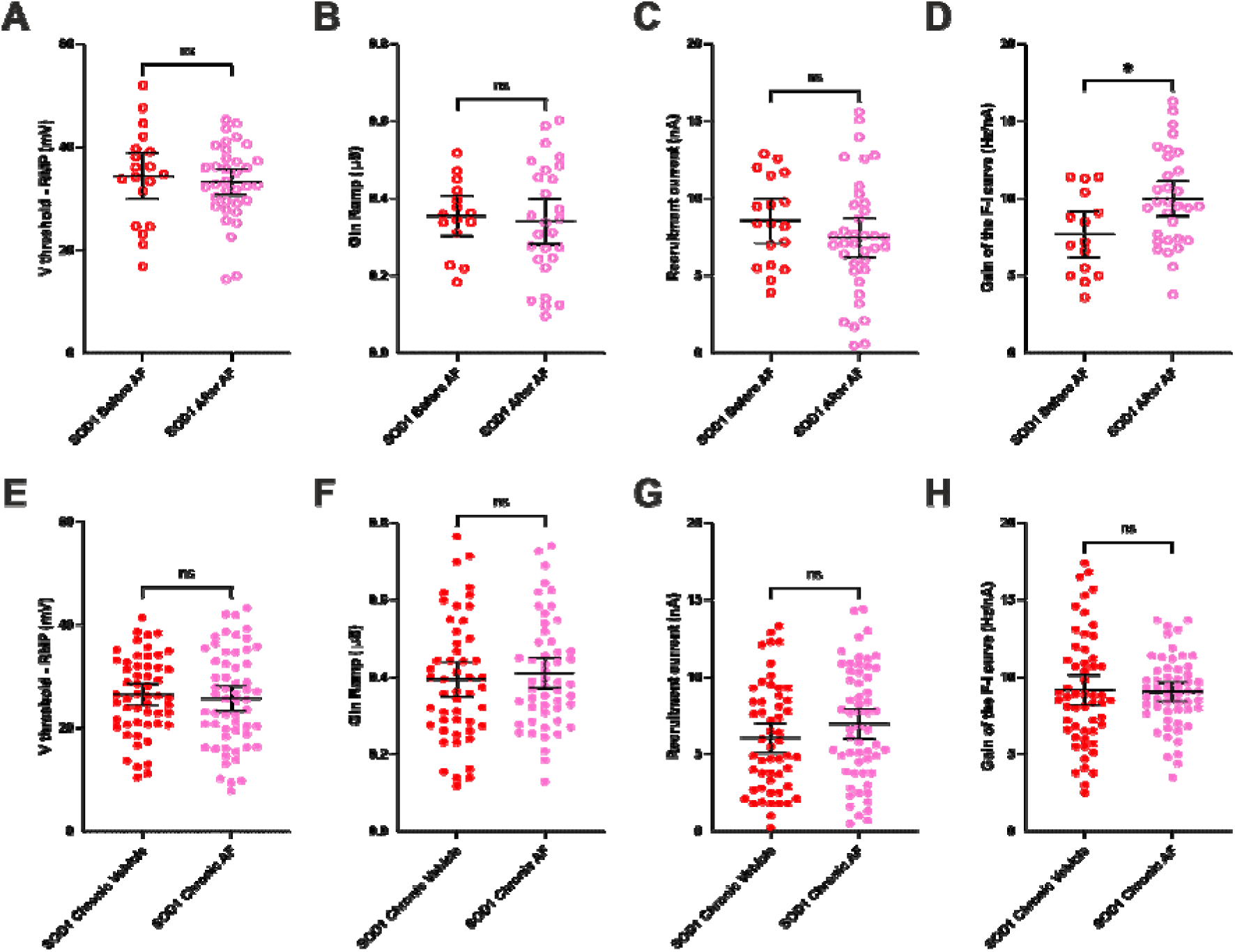
Replication of electrophysiological findings upon acute and prolonged delivery of adrenergic β2/β3 agonists on a different cohort of presymptomatic SOD1 mice. **A-H)** Electrophysiological properties were obtained from slow ramps of current, as in Figure 4 and 7. **A-D)** Effect of the acute treatment on Voltage threshold - resting membrane potential **(A)**, ramp input conductance **(B)**, recruitment current **(C)**, gain of the F-I relationship **(D)**, in MNs from SOD1 mice. E-H) Effect of the chronic treatment on Voltage threshold - resting membrane potential **(E)**, ramp input conductance **(F)**, recruitment current **(G)**, gain of the F-I relationship **(H)**, in MNs from SOD1 mice. In all graphs, each point represents one MN and the mean ± 95% confidence intervals are shown. N = 7 Acute SOD1 mice and N = 11 Chronic SOD1 mice. \**p*<0.05, ns - non significant.

